# From sound to source: Human and model recognition of environmental sounds

**DOI:** 10.64898/2026.03.12.711349

**Authors:** Sagarika Alavilli, Josh H. McDermott

## Abstract

Our ability to recognize sound sources in the world is critical to daily life, but is not well documented or understood in computational terms. We developed a large-scale behavioral benchmark of human environmental sound recognition, built models of sound recognition that operate on the acoustic signal, and used the benchmark to compare models to humans. The behavioral benchmark measured how sound recognition varied across source categories, audio distortions, and concurrent sound sources, all of which influenced recognition performance in humans. Artificial neural network models trained to recognize sounds in multi-source scenes reached near-human accuracy and qualitatively matched human patterns of performance in many conditions. By contrast, classifiers trained to recognize sounds from traditional models of the cochlea and auditory cortex produced worse matches to human performance. Models trained on larger datasets exhibited stronger alignment with both human behavior and brain responses. The results suggest that many aspects of human sound recognition emerge in systems optimized for the problem of real-world recognition.

## Introduction

Environmental sound recognition refers to the process by which listeners identify everyday sounds, such as footsteps, rainfall, or animal calls. This ability allows humans to monitor for events of interest and build a representation of the surrounding environment, even when objects or events are out of sight.

Although it is clear from everyday experience that humans are able to recognize many environmental sounds, our understanding of these recognition abilities remains limited. Studies over the past two decades made initial measurements of human environmental sound recognition abilities [1–5], characterizing the dependence on spectral information [6] and summary statistical properties [7,8], for instance. However, experiments have been limited by the lack of large sets of high-quality sound recordings and by the absence of standardized paradigms for evaluating recognition [9]. As a result, many key questions remain unanswered, including how environmental sound recognition is influenced by concurrent sounds, the extent to which it is robust to environmental sources of variation (background noise, reverberation etc.), and the way in which recognition interacts with selective attention. The growth in availability of large audio datasets raises the possibility of a more thorough characterization of human abilities.

Understanding any perceptual ability also entails being able to build models that can account for that ability in realistic conditions. In this respect environmental sound recognition has also been neglected compared to other areas of audition, with few such candidate models [7]. Candidate models might be found in recent advances in machine hearing that have led to notable improvements in automated environmental sound recognition [10]. For instance, neural network models trained on large-scale datasets [11–13] now perform well enough on classification tasks to be used in real-world applications such as automatic captioning [14]. However, such models have not been systematically evaluated against human behavioral data, and it remains unclear whether they reproduce human recognition performance to any extent.

Recent work in other areas of audition has found that neural network models optimized for auditory tasks tend to exhibit human-like patterns of performance (exhibiting variation in accuracy across conditions that mirrors that seen in humans) despite not being fit to match human performance. This general result has been observed in models of word recognition [15,16], sound localization [17], pitch perception [18], and selective listening [19]. The presumptive explanation of such results is that human perception has been driven towards high performance of ecologically important tasks through a combination of evolution and learning over development, and that our behavioral phenotype is strongly constrained by this implicit optimization process.

Comparisons to task-optimized models provide explanations of human behavior in terms of optimization, while also allowing exploration of the environmental factors that shape human perception. For instance, models of sound localization optimized in virtual environments lacking surface reflections fail to exhibit sound onset dominance in localization behavior, providing evidence that this effect in humans is an adaptation to help robustly localize sounds in the presence of reflections [17]. Conversely, discrepancies between humans and models raise the possibility that additional biological constraints are needed to explain human behavior. For instance, in models trained to estimate fundamental frequency, the inclusion of an initial model stage that simulates high-fidelity cochlear processing is needed to account for human pitch perception [18]. High-fidelity cochlear processing is also necessary to account for aspects of human voice recognition [16]. Being able to build models that reproduce human behavior also opens the door to understanding effects of hearing loss on behavior [20], and to new approaches for understanding prosthetic devices [21].

In this paper we assess whether neural network models optimized for environmental sound recognition exhibit human-like patterns of performance. There were two main goals. The first goal was to set up the infrastructure needed to better characterize human abilities and enable comprehensive human-model comparisons in this domain. We developed a large-scale behavioral benchmark for environmental sound recognition, measuring recognition performance across a range of sound categories, scene sizes, and auditory manipulations. These experiments drew on ideas from previous studies but expanded their scope to provide a larger-scale “fingerprint” of human abilities than had been previously available. The second goal was to use the infrastructure to ask whether task optimization was sufficient to explain human environmental sound recognition abilities, at least in the settings probed by the benchmark. We thus evaluated both in-house and publicly available models on these same benchmarks to assess how well current systems account for patterns of human recognition behavior.

We found that the various conditions in the benchmark produced reliable variation in human recognition performance, providing a behavioral signature that models should account for. Neural network models optimized for environmental sound recognition reproduced this pattern of behavior to a considerable extent, particularly when they were trained on large datasets. By comparison, traditional models based on cochlear and auditory cortical processing stages showed a more limited correspondence with human behavior. The results suggest that optimizing machine learning systems for the problem of sound recognition is a promising approach to developing models of human abilities in this domain.

## Results

### Humans environmental sound recognition benchmark

To benchmark human environmental sound recognition, we conducted three experiments with human participants. In each experiment, participants performed a sound category detection task in which they heard an audio signal and judged whether a particular category was present. We chose this task because it allowed recognition to be queried in scenes with multiple sound sources, without requiring the participant to identify all of the sources. The category query was presented after the scene (except for Experiment 1c, as discussed below) to facilitate comparisons with the models we considered, which did not have a means to direct attention to particular sound categories. For the same reason, participants could not replay the sound. We measured how recognition performance was influenced by the presence of concurrent sounds (Experiment 1) or by audio distortions applied to the sound signal (Experiment 3). Performance was quantified as d’, calculated from hits (defined as “yes” responses when the probed source label corresponded to a source present in the auditory scene) and false alarms (defined as “yes” responses when the probed label did not correspond to any source present in the scene). We will refer to the three experiments as the “EnvAudioEval” benchmark.

### Experiment 1: Effect of scene size on environmental sound recognition

On each trial, participants were presented with an auditory scene – a sound waveform containing the superposition of between one and five sound sources (we henceforth refer to the number of sources as the “scene size”). The scenes were constructed from randomly selected sound categories and the onset of each source was randomly jittered (see Methods). Participants judged whether a specific target category was present within the scene (Figure 1a). We chose an upper limit of five sources based on pilot experiments that showed a range of 1 to 5 was sufficient to produce substantial variation in human recognition performance. Target categories are listed in Figure 1b. For Experiments 1a and 1c, data were collected online using established quality control measures to help ensure participant engagement and headphone usage; Experiment 1b was conducted in person (see Methods).

**Figure 1.**
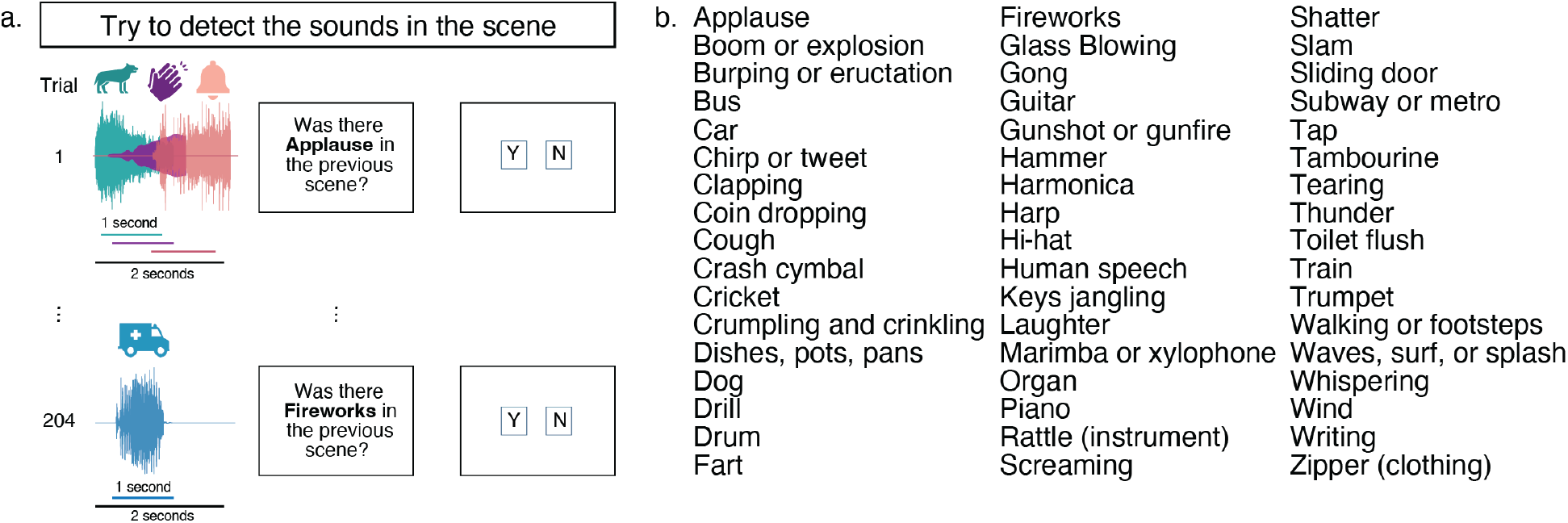
Overview of Experiment 1. A) Experimental design. All sources had jittered onsets and durations of up to 1 second, with a maximum scene length of 2 seconds. Different colors illustrate different sources present within the scene. B) Sound categories used in experiment.

### Experiment 1a: Human recognition declines with scene size

We first analyzed performance as a function of scene size. Performance declined as the number of concurrent sources increased, but remained above chance (corresponding to a d’ value of 0) for scenes of five sources (Figure 2a).

**Figure 2.**
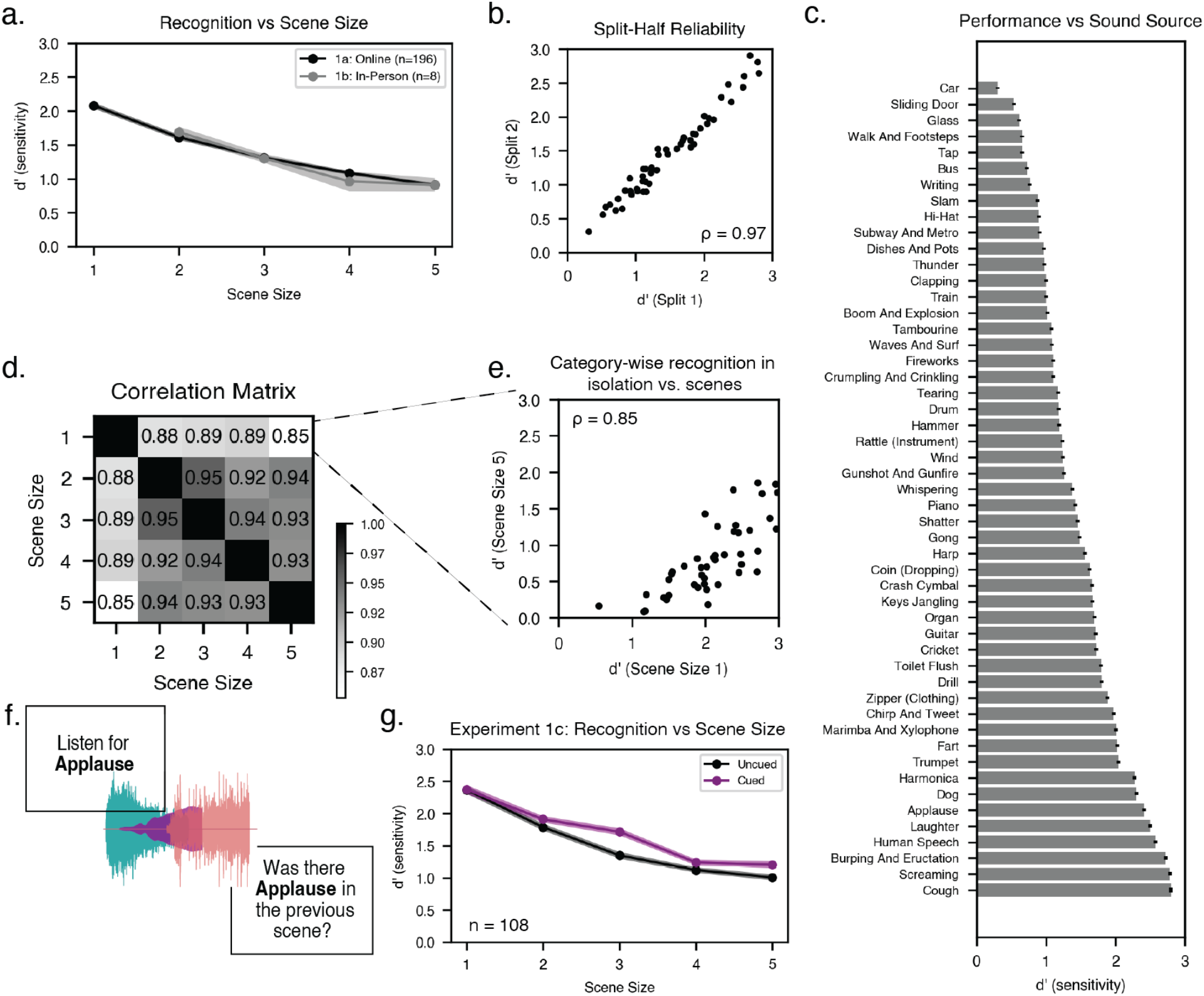
Experiment 1: Human recognition accuracy varies with scene size and sound category. A) Recognition accuracy for different scene sizes. Black line plots results from online experiment (Experiment 1a) and gray line shows results from in-person experiment (Experiment 1b). Error bars plot SEM (may not be visible due to marker size). Chance performance corresponds to a d’ of 0. B) Split-half reliability of accuracy for different sound categories, averaged across scene sizes (data from Experiment 1a). C) Recognition accuracy for each sound category in Experiment 1a. Sound categories are ordered by increasing d’. Error bars plot SEM. D) Correlations between category-wise recognition performance (d’) for different scene sizes in Experiment 1a. E) Comparison of recognition accuracy for each category when presented in isolation (scene size of 1) vs. in 5-sound scenes (data from Experiment 1a). F) Example trial from the cued condition of Experiment 1c (uncued condition had the same trial format as Experiment 1a). G) Results of Experiment 1c: recognition accuracy vs. scene size for cued (purple) and uncued (black) conditions.

### Experiment 1a: Humans show reliable variation in recognition across categories

We also analyzed recognition performance in Experiment 1a as a function of the probed sound category, averaging across scene sizes. This variation was highly reliable across splits of participants (split-half reliability of d′ across sound categories: ρ = 0.97, p < 0.001; Figure 2b). Human performance varied substantially by category: some categories (e.g., coughing) were highly recognizable, while others showed consistently lower recognition rates (e.g., car; Figure 2c).

The variation in recognition performance across categories could in principle reflect both the distinctiveness of a sound category relative to the other categories as well as factors that might be specific to detecting a sound when presented in a scene with other sounds. For instance, when presented with other sounds, some sound categories might be less likely to be masked than others. Masking is typically driven by overlap in peripheral or central representational channels, and thus could be less likely in sound categories with unusual distributions of spectral energy or other features. Other potential scene-specific effects could include biases for more familiar sound categories.

To probe for scene-specific influences on recognition, we measured the correlation between recognition performance by sound category for each scene size. If the factors that determine sound recognizability differ between isolated presentation and multi-source scenes, then recognizability measured in silence should show a reduced correlation with recognizability across other scene sizes. We calculated the correlation between category-wise performance for each scene size (Figure 2d). The sample size was large enough that these correlations were themselves fairly reliable (split-half reliability of the off-diagonal elements of the correlation matrix was 0.500). Category-wise performance was strongly correlated across scene sizes, but correlations between recognizability for isolated sounds (scene size = 1) and recognizability for other scene sizes were lower than analogous correlations between pairs of scene sizes greater than one. For instance, recognizability in a scene size of 2 was more strongly correlated with recognizability in other multi-source scene sizes (e. g. scene size of 5: ρ = 0.94) than with that in a scene size of 1 (ρ = 0.88). All four of the correlations involving a scene size of 1 were lower than the correlations not involving a scene size of 1, which is unlikely to happen by chance (p=.0048, permutation test). These results indicate that although some of the variance in recognizability across categories reflects category distinctiveness, other scene-level factors also play some role in determining a sound’s recognizability in a larger scene.

We note that the factors that underlie a sound’s recognizability in isolation might also indirectly determine its recognizability in auditory scenes with other sounds. For instance, if a sound category is acoustically distinct from other categories (making it easier to recognize in isolation), it might also be less prone to masking due to decreased spectrotemporal overlap with other sounds. One thus might expect recognizability in isolation to be correlated with recognizability in larger scenes, even if the mechanisms constraining recognizability in the two cases are distinct.

### Experiment 1b: In-person replication

To ensure that human performance was not being underestimated as a consequence of limitations of the online experiment setting, we ran a second experiment with in-person participants who heard sounds played from a loudspeaker. The experiment presented scenes of 2-5 sources, with the same task as Experiment 1a. As shown in Fig. 2a, performance levels were similar in the in-person setting as in the online setting, consistent with prior studies showing that online experiments generally yield similar results to in-person experiments provided measures are taken to ensure reasonable-quality sound presentation and participant compliance [15,22–28].

### Experiment 1c: Cueing manipulation

To assess whether human performance was limited by the sound category query appearing after the scene, we conducted an additional online experiment in which listeners were cued to the queried sound category before the scene on half of the trials (Figure 2f). The experiment was otherwise identical to Experiment 1a. As shown in Fig. 2g, performance was similar with and without the category cue for single sources, but diverged for larger scenes, with better performance in the cued condition. Specifically, there was a main effect of cueing (F(1,107)=17.09, p<0.001), but no significant difference between the cued and uncued conditions for single sources (t(107)=0.52, p=0.60). This result is consistent with the idea that directed attention (in this case, to a category of sound) aids perception in multi-source scenes, but has less of an effect when only one source is present.

### Experiment 2: Sound category confusions

To better understand the potential confusability of different sound categories, we ran a second online experiment using the same task as Experiment 1a but featuring exclusively single sources. To systematically characterize confusability across categories, we designed the experiment such that every off-diagonal cell in the confusion matrix was presented the same number of times across participants, ensuring uniform coverage of potential category confusions.

Despite the fairly low number of trials contributing to each off-diagonal cell (eight in total), the confusion matrix exhibited reliable structure (Fig. 3A; split-half reliability of the off-diagonal elements was r=0.52, p=0.001). Moreover, inspection of the matrix reveals that some categories were frequently confused. For instance, participants judged the “Boom and explosion” category to be present every time for the sound of fireworks. Similarly, train sounds were confused for “subway and metro” and keys jangling were confused as “coins dropping”. These results indicate that some pairs of categories have highly overlapping distributions of sounds, such that there is a limit to how well any recognition system should be expected to perform.

**Figure 3.**
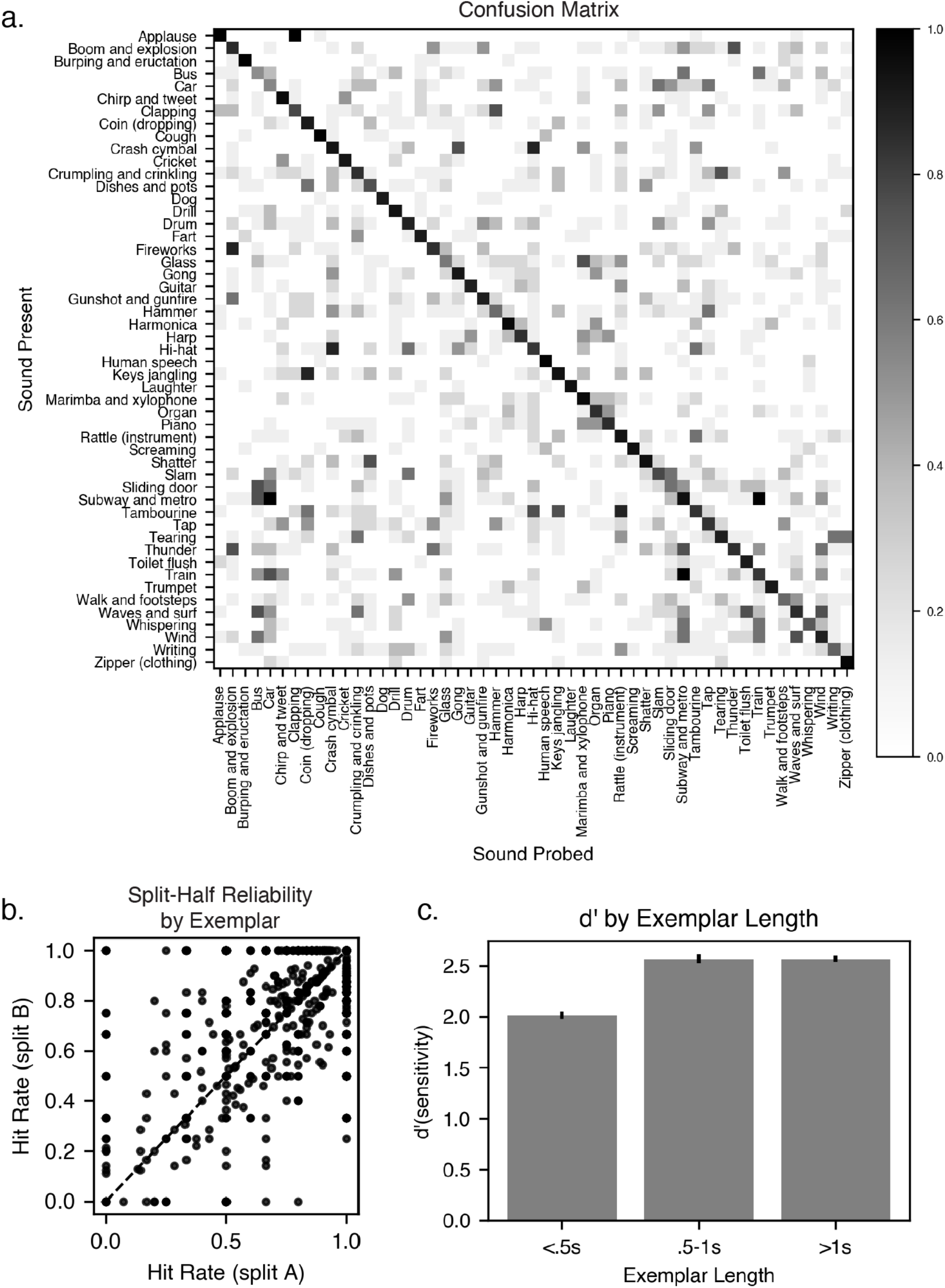
Experiment 2: Human sound category confusions. A) Confusion matrix showing the proportion of “yes” responses for pairings of sound category queries (horizontal axis) and sound category presented (vertical axis). B) Split-half reliability of hit-rate of exemplars. C) Recognition accuracy for different sound lengths. Error bars plot SEM.

This experiment also permitted an analysis of whether particular exemplars from a category were consistently better recognized than others. We analyzed all stimulus exemplars that were presented and probed with their true label at least four times, splitting the trials into two halves, and calculating the split-half reliability of the hit rates in each half. As can be seen in Fig. 3B, these hit rates were fairly reliable even given the modest number of trials involved (r=0.66, p<.001). This result indicates that there is meaningful structure at the exemplar level, which might be useful for comparing humans and models. It also suggests that some exemplars are not reliably detected as their labeled category (though most are detected most of the time).

One other factor that could have limited human performance is the truncation of sounds to 1s in length. To address this issue, we analyzed performance as a function of the source sound length, leveraging the fact that many of the original sounds were less than 1s in length. As shown in Fig. 3c, performance was worst for the shortest sounds (i.e., those that were not truncated), and improved with sound duration (F(2, 398)=72.68, p < 0.01, showing a main effect of duration). These results do not exclude the possibility that performance could be somewhat better if the longer sounds were not truncated, but there is no evidence that the truncation somehow interferes with recognition.

### Experiment 3: Effect of sound distortions on environmental sound recognition

Sound distortions, in which particular aspects of a sound are altered or degraded, are often used to probe the acoustic features that humans rely on for recognition [6,29,30]. The second component of our benchmark involved measuring the effect of a large set of distortions on human recognition, as this forms an additional aspect of the “fingerprint” of human abilities that can be compared to models. The task was the same as in Experiment 1a, but with scenes consisting of a single source to which a distortion had been applied.

We assembled a set of distortions commonly used in studies of auditory perception, including local time reversal [31], time dilation and compression, reverberation [32,33] (varying either reverberation time or the direct-to-reverberant ratio [33]), clipping [34], noise vocoding [29], bandpass filtering (varying either the bandwidth or center frequency of the pass-band) [6,35], lowpass filtering, highpass filtering, and modulation filtering [30] (lowpass in either the spectral or temporal domains). Each distortion was applied at several magnitudes that distorted the original audio to different extents.

Each type of distortion tests the dependence of recognition on a different aspect of sound structure. Audio frequency manipulations (high-pass, low-pass, and band-pass filtering) probe the degree to which recognition depends on specific frequency regions and/or is invariant to the “channel” through which sound is heard. Time compression and dilation test invariance to the rate of audio events [36,37]. Local time reversal and temporal modulation filtering disrupt the structure of signals at different timescales, isolating the role of fine-vs. coarse-grained temporal structure [38,39]. Reverberation also disrupts temporal structure, but in a way that occurs frequently in everyday listening. Spectral modulation distortions (spectral modulation filtering and noise vocoding) test the resolution of spectral detail needed for recognition. Peak clipping tests tolerance for harmonic distortion and reductions of dynamic range. Spectrograms in Figure 4a illustrate the effects of a subset of distortions on three example sounds.

**Figure 4.**
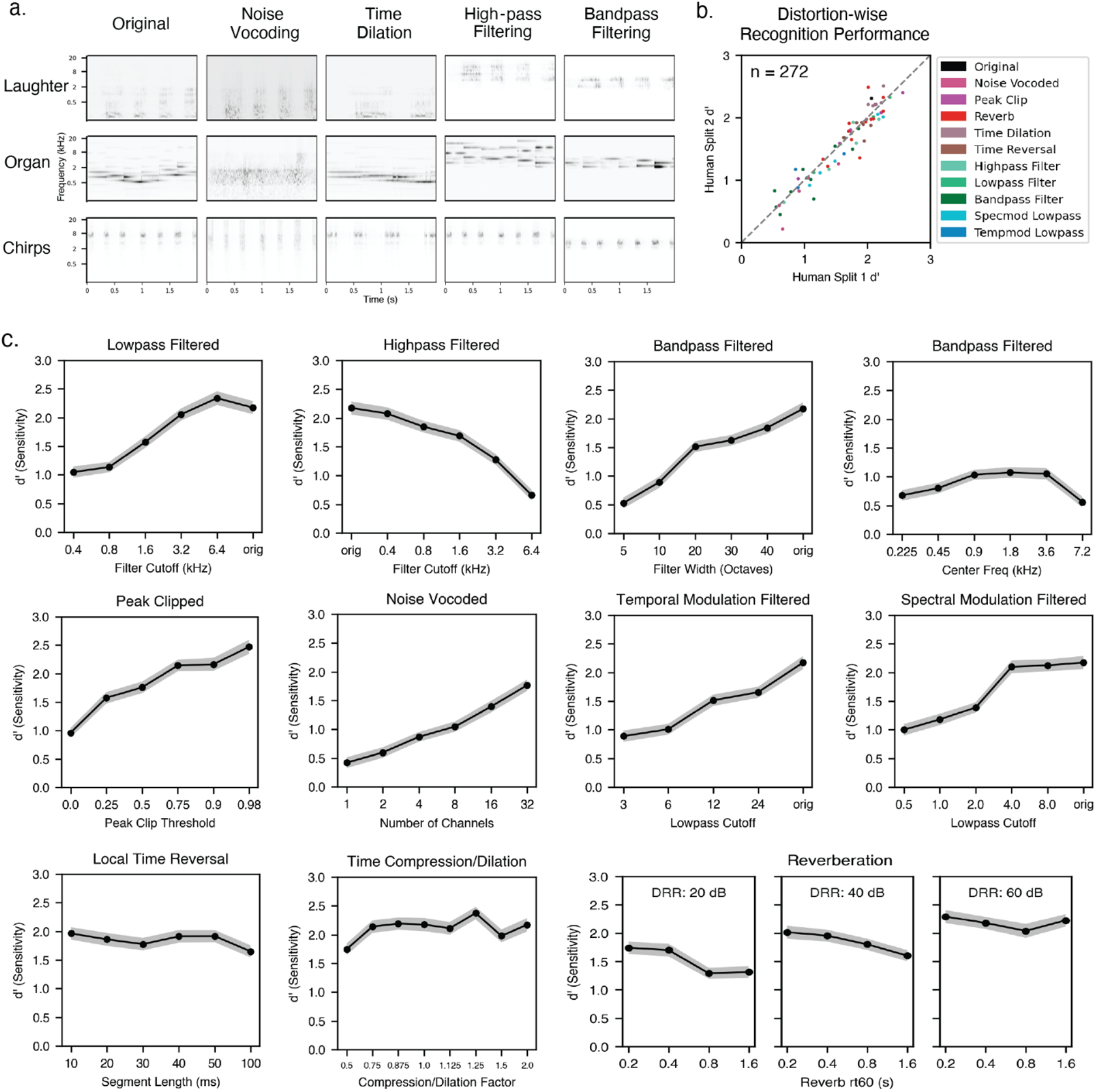
Experiment 3: Effect of distortions on human environmental sound recognition. A) Spectrograms of three example sounds processed with each of a subset of the tested distortions. Sounds were renormalized after the application of the distortions, which is why frequencies are visible in the audio filtering conditions that are not visible in the original. B) Split-half reliability of recognition across distortion conditions. C) Human recognition performance as a function of distortion level for all distortion types. Error bars plot SEM.

### Humans show reliable variation in recognition across sound distortions

Human recognition exhibited consistent variation across distortion types and levels (Figure 4b; split-half reliability was high (ρ=0.95, p < 0.001). Some distortions markedly impaired recognition, while others had little effect (Figure 4c). Performance declined most substantially when frequency information was eliminated (e. g. high-pass, low-pass, or band-pass filtering), whereas temporal manipulations such as time dilation and local time reversal had smaller effects on recognition performance (Figure 4c). Deleterious effects of audio filtering, and the crossover point between the lowpass and highpass psychometric functions, are consistent with previous findings in both sound recognition and speech perception [6,29,30,34,40]. In the noise-vocoded condition, performance (d’) remained above chance even when spectral resolution was reduced to a single channel, indicating that temporal-envelope cues alone are sufficient to support some degree of recognition in the absence of any spectral detail. The robustness of recognition to local time reversal further suggests that the fine-grained temporal envelope contributes little to environmental sound identification. Recognition was fairly robust to reverberation, potentially due to its regular presence in everyday environments [33].

We observed some notable differences between the effects of particular distortions on environmental sound recognition compared to speech recognition. The two clearest differences were obtained with time compression/dilation and local time reversal. Both of these manipulations had relatively little effect on environmental sound recognition, contrasting with strong previously documented effects on speech [31]. This difference likely indicates that fine-grained temporal structure is less diagnostic of environmental sounds than it is for speech. Consistent with this idea, the most lowpass temporal modulation filter (cutoff of 3 Hz) left environmental sound recognition well above chance (d’ of ∼1), whereas it renders speech almost fully unintelligible [30]. By contrast, the degradations of spectral structure (peak clipping, noise vocoding, and spectral modulation filtering) have sizeable effects on speech recognition [29,30,34,41], and also had large effects on our task. The effects of audio filtering were also qualitatively similar to those previously observed for speech recognition [42,43], potentially indicating a domain-general ability to discount “channel” effects.

### Modeling Overview

We evaluated computational models falling into three categories. First, to assess whether traditional models with hand-engineered features could explain human behavior, we evaluated baseline models consisting of optimized linear classifiers operating on standard biologically inspired filterbanks. Second, we trained a set of artificial neural network models from scratch to perform the task. Model architectures were adapted from those that previously produced strong performance on other auditory tasks (see Methods for details). Third, we tested models that incorporated alternative architectures and/or benefited from large-scale pretraining on other audio datasets. These included convolutional and transformer-based models selected based on their performance on the AudioSet benchmark [44]. AudioSet is currently the largest labeled dataset for environmental sound recognition, and the sound categories used in the EnvAudioScene dataset, and in the EnvAudioEval experimental benchmark, were a subset of the classes in AudioSet (but with nonoverlapping sets of sound exemplars), specifically those used in the subsequent GISE-51 dataset [45]. In all cases the models were trained or fine-tuned on the same dataset of scenes constructed from recorded environmental sounds, see Methods). All models had independent classifiers for each sound category, and were trained or fine-tuned with a separate binary cross-entropy loss for each category. Figure 5 shows schematics of each of the model architectures.

**Figure 5.**
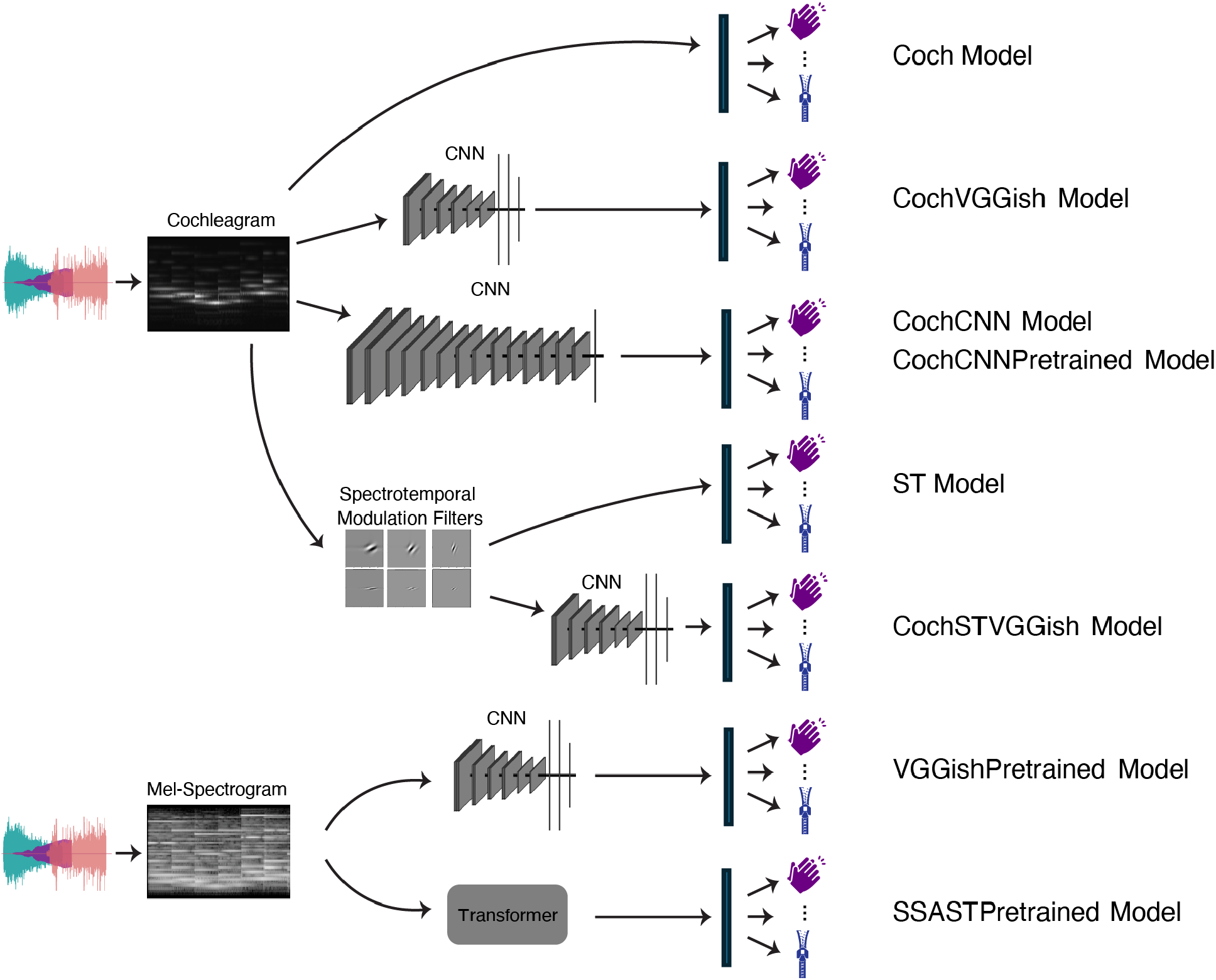
Model architectures. Audio waveforms were transformed to either a cochleagram (a time-frequency decomposition generated using a simulation of the human cochlea) or a mel-spectrogram (a spectrogram with frequency bins clustered to approximate the frequency resolution of human hearing).

### Training data

Models were trained of fine-tuned on a dataset of scenes constructed from sounds drawn from the GISE-51 [45] dataset. GISE-51 consists of recordings of isolated sounds in 51 categories that are a subset of the categories in the FSD-50k dataset [12]. We generated scenes of between 1 and 5 randomly selected sound sources, rendered with realistic reverberation. We note that this training data generation procedure omits statistical regularities in the sources that co-occur in naturally occurring soundscapes, which could produce differences in the recognition strategies of the model when compared to humans. Each scene had a category label for each source. We refer to this as the EnvAudioScene dataset (1.5 million scenes constructed from 16,357 sound clips in GISE-51). EnvAudioScene is smaller than the AudioSet dataset, but is more controlled, as the scenes were synthesized from clean source recordings (AudioSet consists of soundtracks from YouTube videos).

### Baseline models

The cochleagram model (Coch) applied a cochlear filterbank to the input waveform (to approximate the human auditory periphery) followed by a linear classifier (Figure 4a). The spectrotemporal model (ST) used a spectrotemporal filterbank often used to approximate primary auditory cortical processing [46–48]. The model consisted of the cochlear stage from the cochleagram model, followed by a spectrotemporal filterbank, followed by a linear classifier (Figure 4b). Both models were trained on the EnvAudioScene dataset.

### In-house models

The in-house model was based on the top-performing architecture from a previous publication [16]. The architecture consisted of the fixed cochlear stage used in the baseline models followed by a convolutional neural network (CNN). We tested a variant of this model trained just on EnvAudioScene (CochCNN) as well as a variant that was pretrained on AudioSet (CochCNNPretrained) and then fine-tuned on EnvAudioScene. We also tested a variant that incorporated the spectrotemporal filterbank from the ST baseline model prior to the CNN (CochSTVGGish). We used the VGGish [11] architecture as the CNN backbone for this model since its input dimensions most closely approximated the output dimensions of the ST model. The dimensions of the VGGish input layer were updated to accommodate the increased number of frequency channels and input length (see Methods).

### External models

We evaluated two externally developed architectures: VGGish and SSAST. The VGGish model [11] is a CNN trained on AudioSet with mel-spectrogram inputs. We tested the original model after fine-tuning on EnvAudioScene (VGGishPretrained), as well as a model with the VGGish architecture applied to cochleagram input and trained from scratch on EnvAudioScene (CochVGGish). The SSAST model [13] is a self-supervised version of the Audio Spectrogram Transformer [49]. SSAST was pretrained on Librispeech [50] and AudioSet [44] using a self-supervised learning objective (SSAST). We then fine-tuned this model on EnvAudioScene prior to evaluation.

### Models replicate human patterns of performance for sound recognition

We evaluated model performance on the same task performed by human participants, using each of the two human benchmark experiments. We used the same stimuli as in the human experiments. Model performance was calculated as the area under the receiver operating characteristic curve (AUC), converted to d’ for comparison with humans. For both experiments, we examined whether models replicated overall levels of human performance, as well as the pattern of performance across conditions.

### Human and model recognition degrades similarly in multi-source scenes

All models qualitatively replicated the dependence of human recognition performance on scene size, with recognition accuracy highest for single-source scenes and declining as the number of sources increased (main effect of scene size for all models: F(4, 250) > 13.22, p < 0.01 in all cases). However, the fully optimized CNN-based models most closely matched human levels of performance across scene sizes (root-mean-square error between model and human results: Coch_Model: 1.33, CI=(1.30, 1.36), ST_Model: 0.91, CI=(0.88, 0.94), CochCNN_Model: 0.44, CI=(0.42, 0.46), CochVGGish_Model: 0.62, CI=(0.59, 0.64), CochSTVGGish_Model: 0.62, CI=(0.60, 0.65), CochCNNPretrained_Model: 0.42, CI=(0.40, 0.43), VGGishPretrained_Model: 0.39, CI=(0.37, 0.41), SSASTPretrained_Model: 0.38, CI=(0.37, 0.39)), while the baseline models underperformed relative to human listeners (Figure 6a). The models pretrained on large amounts of data (CochCNNPretrained, VGGishPretrained, and SSASTPretrained) exhibited the best quantitative match to human performance.

**Figure 6.**
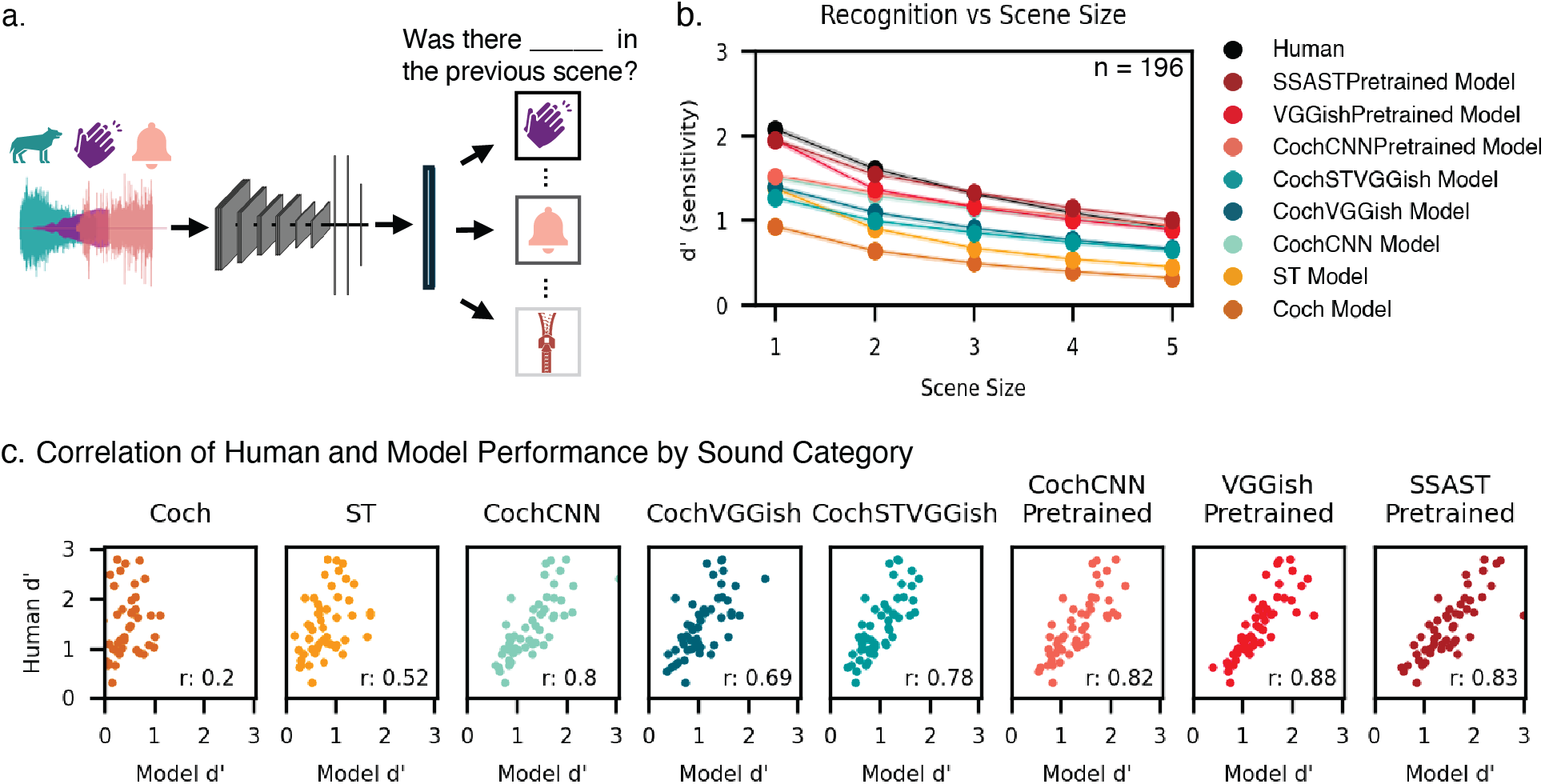
Model recognition of sound sources in auditory scenes (Experiment 1a). A) Schematic depicting the evaluation pipeline in which models were tested on the identical auditory scenes presented to human participants. B) Model recognition performance vs scene size. C) Comparison of human and model recognition accuracy. Model d′ values are plotted on the x-axis and human d′ values are plotted on the y-axis of each plot; each point represents one of the 51 sound categories.

### Pretrained models best replicate human performance by sound category

Figure 6b compares human and model performance for individual sound categories. The baseline models underperformed relative to humans. By contrast, the optimized neural network models more closely matched human performance, and captured much of the variance in human recognition across categories (Figure 6b; human-model correlations ranged from.69 to.88). The best performing model (SSASTPretrained) was not significantly worse than humans (paired t-test, t(50) =-0.91, p = 0.37); all other models significantly underperformed humans (paired t-test, p<0.04 in all cases). Models pretrained on large datasets and subsequently fine-tuned on GISE-51 (CochCNNPretrained, VGGishPretrained, and SSASTPretrained) yielded the highest human-model correlation, suggesting that larger (and potentially more diverse) training data yields better matches to human behavior (Figure 6c). However, the human-model correlation remained below the human noise ceiling even for the most similar models (the reliability of category-wise performance for the human sample was ρ = 0.99, estimated by Spearman-Brown correcting the split-half reliability measured in Fig. 2B). Some of the differences between model and human patterns of performance may reflect the distributions of the sound categories in the training data. All categories were presented with equal probability during model training, which is unlikely to reflect the distribution of sound categories in real-world human exposure.

### Task-optimized models replicate human patterns of confusions to some extent

To assess whether models replicate the pattern of human sound category confusions, we computed the correlation between human and model confusion matrices using all off-diagonal elements (Figure 7a). All optimized models exhibited positive correlations with humans, but the task-optimized models were again substantially more similar to humans than the baseline models (Figure 7b; Coch_Model: ρ = 0.19; ST_Model: ρ = 0.19; CochCNN_Model: ρ = 0.31; CochVGGish_Model: ρ = 0.31; CochSTVGGish_Model: ρ = 0.31; CochCNNPretrained_Model: ρ = 0.29; VGGishPretrained_Model: ρ = 0.36; SSASTPretrained_Model: ρ = 0.33; p < 0.001 in all cases). However, for all models, the human-model similarity was well below the noise ceiling set by the human confusion reliability (dashed line in Figure 7b), indicating that variance in category confusions is not fully captured by current models.

**Figure 7.**
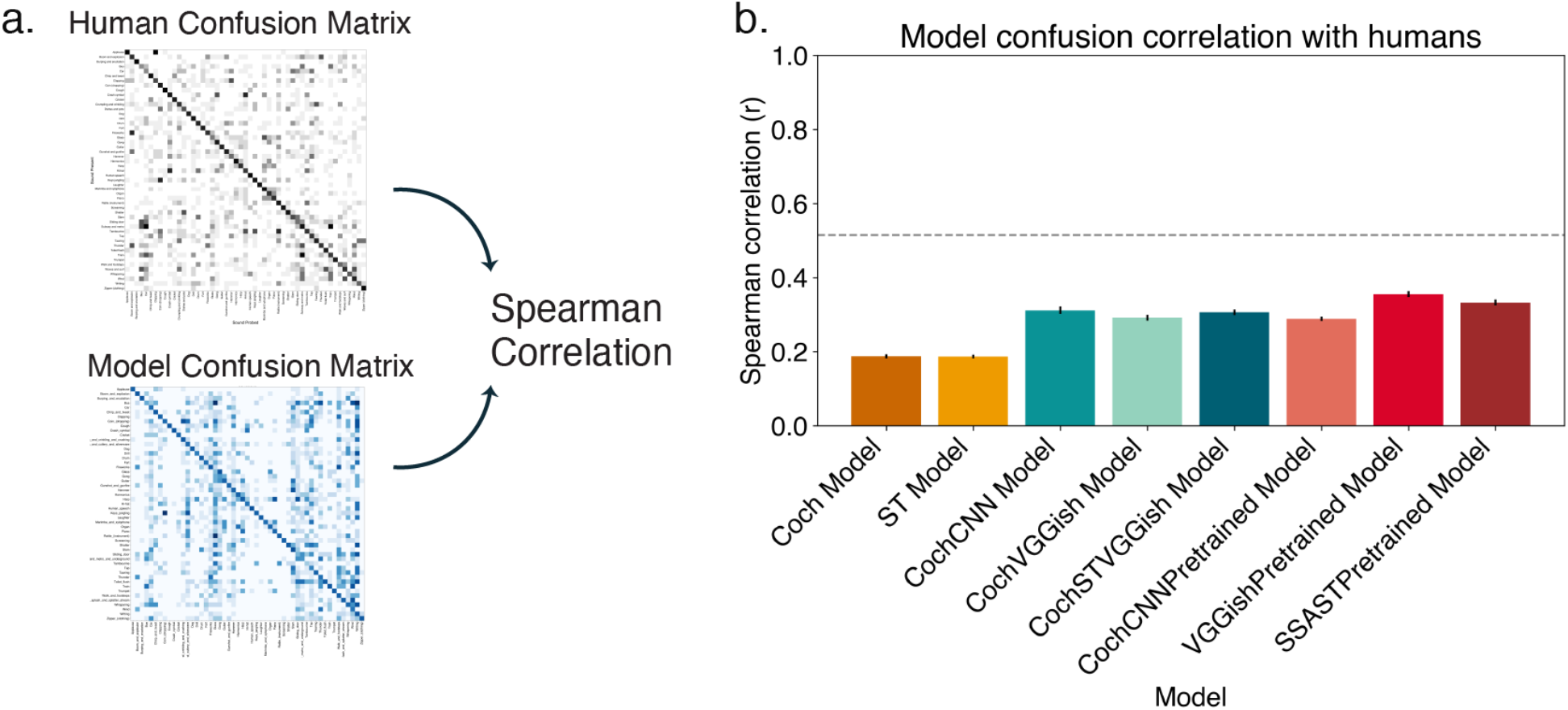
Human-model confusion matrix correlations (Experiment 2). A) Similarity between human and model confusion patterns was quantified by correlating off-diagonal confusion matrix elements. B) Spearman correlation values for each model. Error bars represent standard error of the mean (SEM) estimated via bootstrapping over trials. Gray line indicates human noise ceiling.

### Task-optimized models replicate human exemplar-level variance to some extent

We additionally examined whether models replicate human performance at the level of individual sound exemplars by correlating human and model hit rates for the individual stimulus exemplars analyzed in Figure 3b. All models exhibited significant positive correlations with human exemplar-level hit rates (Coch_Model: ρ = 0.07, p = 0.002; ST_Model: ρ = 0.28, p < 0.001; CochCNN_Model: ρ = 0.39, p < 0.001; CochVGGish_Model: ρ = 0.33, p < 0.001; CochSTVGGish_Model: ρ = 0.38, p < 0.001; CochCNNPretrained_Model: ρ = 0.40, p < 0.001; VGGishPretrained_Model: ρ = 0.48, p < 0.001; SSASTPretrained_Model: ρ = 0.43, p < 0.001), with the pretrained models yielding the strongest correspondence to human behavior. As for the category confusion analysis, the human-model exemplar-level similarity remained below the human noise ceiling in all cases. We note that this analysis is limited by the fact that exemplars that contribute to this analysis tend to be those from sound categories with fewer unique exemplars in the test dataset (as these are the exemplars that tend to be presented more than once in the experiment). These categories are also those that have fewer unique exemplars in the training data. As a result, the sound categories most heavily weighted by this measure are also those for which models are plausibly undertrained, such that the exemplar-level correlations reported here may not be fully representative of model-human correspondence across the broader range of sound categories.

### Pretrained models best reproduce human performance by sound distortion

Figure 8a&b compare human and model performance across sound distortions (color denotes the distortion type and each point represents a specific distortion at a specific level). All models tended to capture the qualitative effect of each distortion on human performance, but humans were overall more robust than the models (every model was significantly worse than humans as evaluated with a paired t-test test comparing the performance for each distortion type and level; p<0.001 in all cases). Both baseline models substantially underperformed relative to humans. All models captured some variance in human recognition performance across distortion levels (with a positive correlation between human and model performance across conditions: r = 0.59 to r = 0.93). We note that the models were not explicitly trained on any of the distortions (with the exception of reverberation, which was present in the training data for the purpose of generating realistic scenes, and with the acknowledgement that it is difficult to say whether these distortions were fully absent from the AudioSet training data that the pretrained models were trained on).

**Figure 8.**
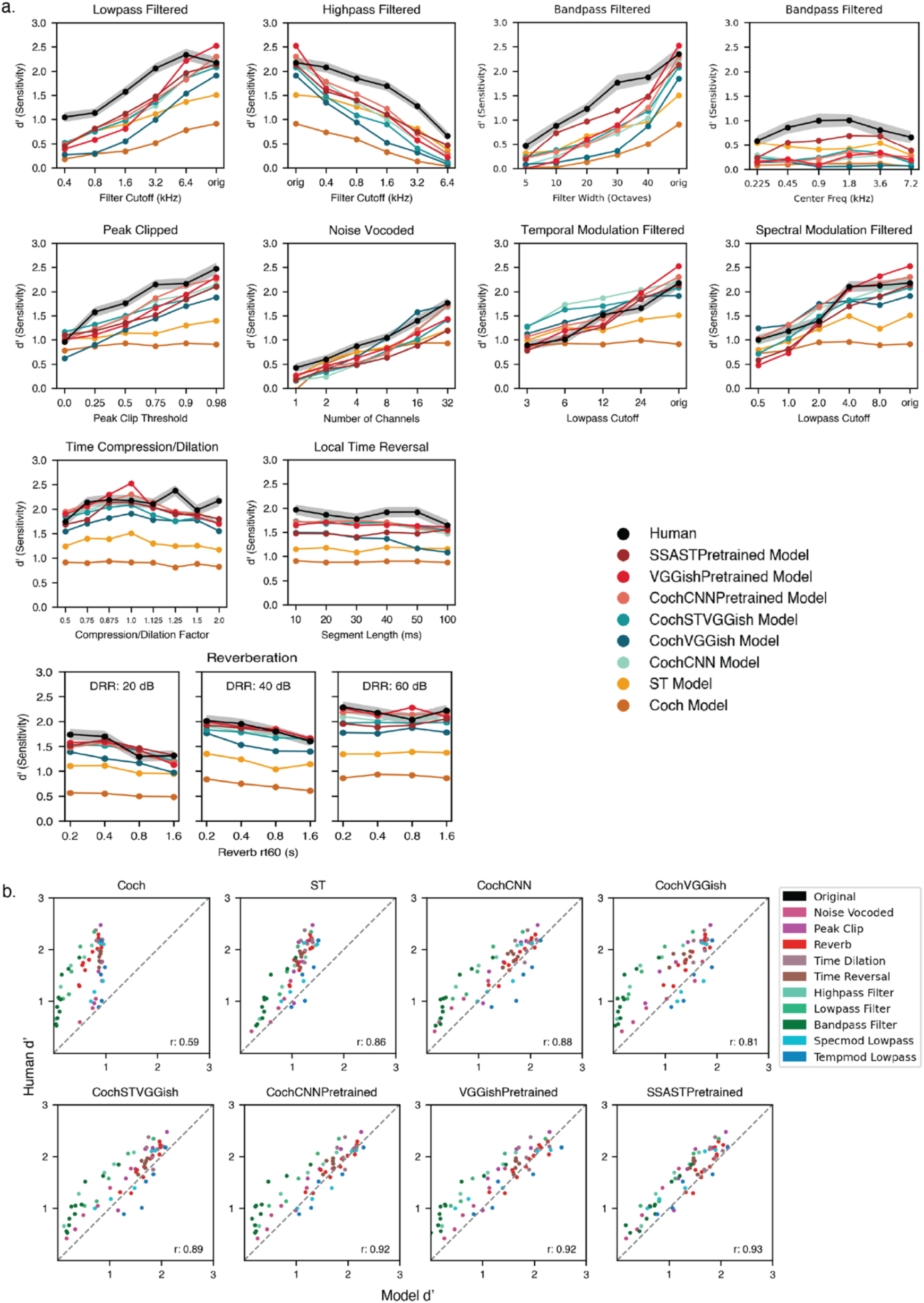
Model recognition of distorted sound sources (Experiment 3). A) Human and model performance for each type of sound distortion. Error bars on human results plot SEM. Error bars on model results are omitted to avoid clutter. B) Human-model correlation of performance across sound distortions. Each panel plots results for a different model, and each dot plots results for one level of one distortion.

CNN-based models—both CochCNN and CochVGGish variants trained solely on GISE-51— approached human-level performance but were less robust for some distortion types, particularly audio filtering (plotted in green in Figure 8a). The inclusion of the spectrotemporal filter stage resulted in somewhat greater robustness to such distortions (significantly better performance for CochSTVGGish than CochVGGish, t(67)=6.25, p<0.001, via a paired t-test comparing performance for each distortion type and level). The models pre-trained on large datasets (CochCNNPretrained, VGGishPretrained, and SSASTPretrained) again yielded the best match with humans (as measured by the correlation between human and model performance). Of these three models, SSASTPretrained, by virtue of self-supervision, was trained with the most data (and had the largest architecture), and also was most robust overall, though the difference between SSASTPretrained and VGGishPretrained was not significant based on a paired t-test across distortions (t(67)=1.09, p=0.28). This result suggests that one of the primary limitations of the other models may be the restricted diversity of the EnvAudioScene training data (inherited from the GISE-51 source recordings). Exposure to the broader acoustic variability in AudioSet, and to the even larger datasets that can be leveraged with self-supervision, appears to improve model generalization, yielding better and more human-like performance.

All models particularly underperformed humans on the audio filtering distortions. By comparison, humans and models were similarly robust to modulation filtering. This result suggests that some aspect of the training and/or architecture leaves these models more dependent on the spectrum compared to human listeners. For instance, the training data plausibly has less spectral variation due to environmental filtering that the experience of a typical human, who hears sounds through walls, windows and other surfaces, and with greater variation in source-listener distance than was present in our training data. Whether or not human robustness is dependent on such experience remains a matter of speculation.

Fig. 9 shows summary bar graphs comparing human-model similarity on the two benchmark experiments. The trained models come closest to the reliability ceiling, computed as the geometric mean of human and model split-half reliability, for the correlation between human and model performance (figure 9a and 9b). Trained models also exhibit lower root-mean-squared error between human and model performance as compared to the baseline models (Figure 9c and 9d).

**Figure 9.**
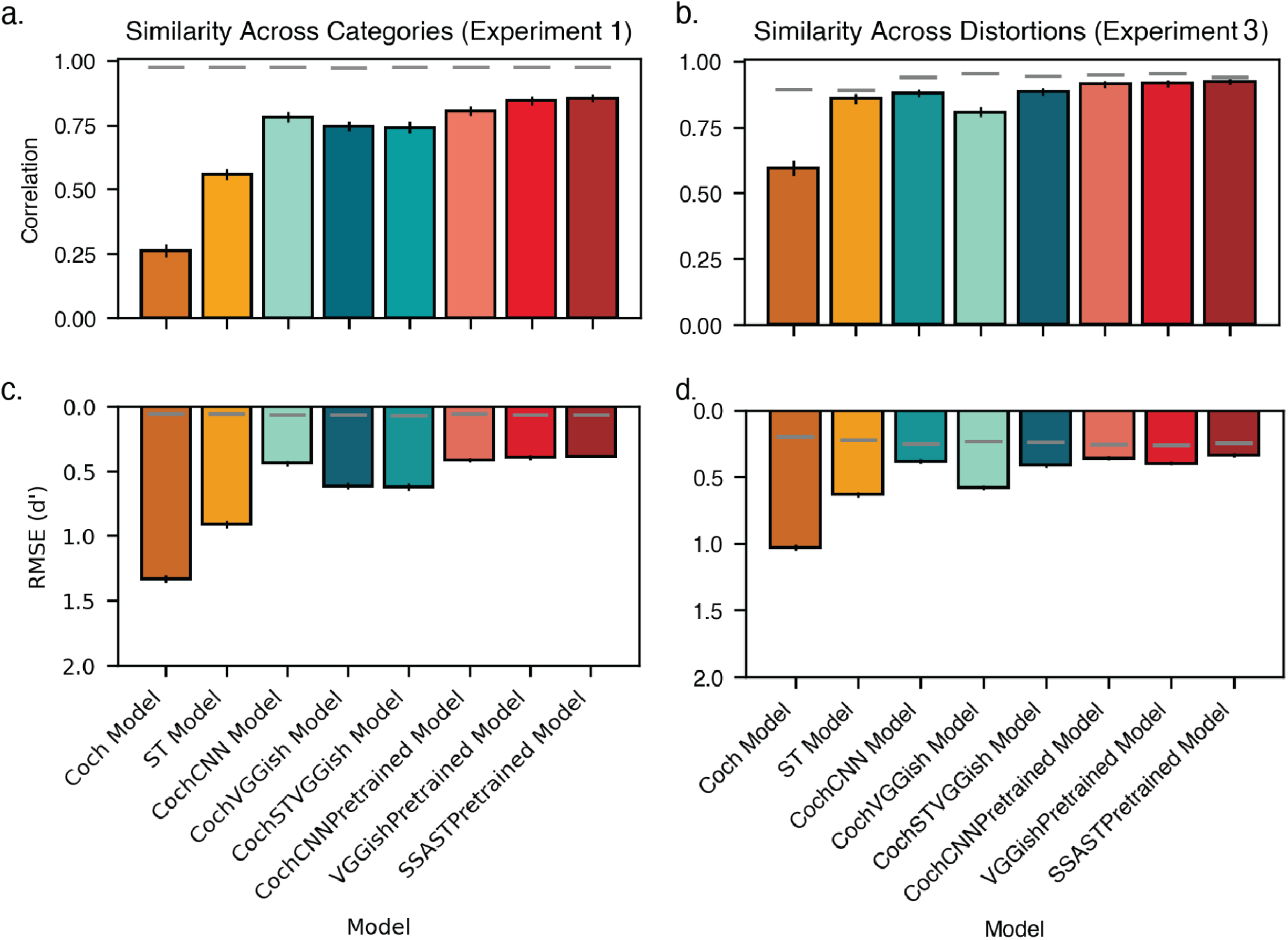
A) Summary plot of correlations between human and model performance (d’) across sound categories (Experiment 1a). Each bar plots this correlation for one of the models. Error bars plot SEM. Gray line indicates the noise ceiling for human-model similarity, calculated using the reliability of the human and model results. B) Summary plot of correlations between human and model performance (d’) across sound distortions (Experiment 3). Gray line indicates noise ceiling (which can differ across models due to variation in the reliability of model performance). Error bars plot SEM. C) Same as A, but plotting root-mean-squared error (RMSE) between human and model category-wise performance. The y-axis is inverted to align the orientation with the correlation-based similarity metrics in A and B. Error bars plot SEM. Gray line indicates the noise ceiling for human-model performance, calculated using the reliability of the human and model results. D) Same as B, but plotting RMSE between human and model performance across sound distortions. Error bars plot SEM. Gray line indicates noise ceiling (which can differ across models due to variation in the reliability of model performance).

### Better models of behavior show higher brain alignment

To evaluate whether models that better replicate human behavior also better reproduce human brain representations, we measured model-brain similarity using the standard metrics of regression-based predictivity and RSA-based representational similarity applied to auditory cortical fMRI responses (Figure 10a&b). The fMRI responses were measured from humans listening to natural sounds [51] (performing an intensity discrimination task intended to maintain attention on the sounds). We replicated the analysis procedures of Tuckute, Feather et al. (2023) [52], who conducted a model-brain similarity analysis of a large set of audio models available at that time.

**Figure 10.**
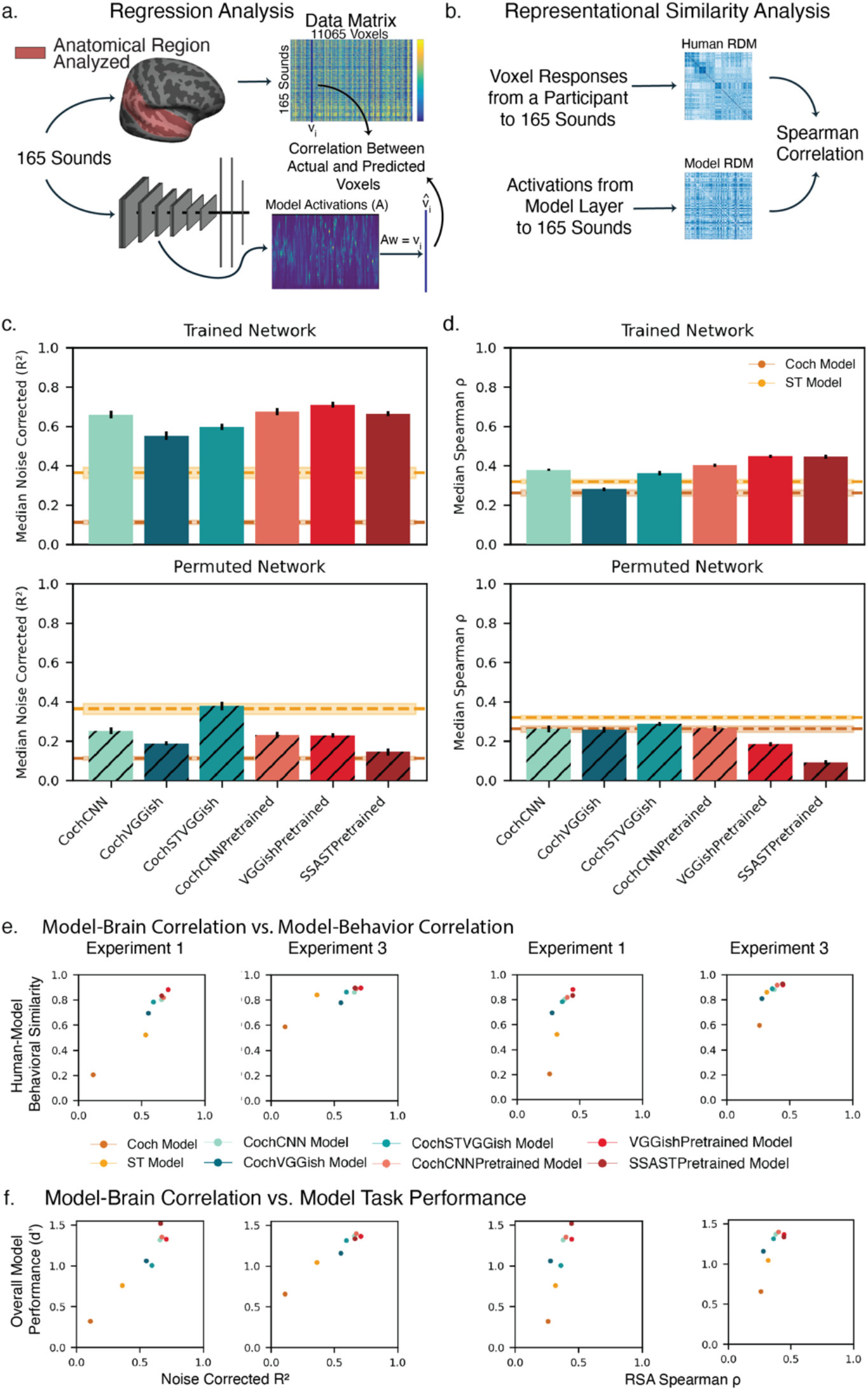
Comparison of model and brain representations of natural sounds. A) Schematic of regression analysis. Voxel-wise brain activity was measured for a set of 165 sounds. For the same stimulus set, time-averaged activations were extracted from a given stage of a computational model. These model-derived features were then used as predictors in a regression framework to account for variance in the fMRI voxel responses. B) Schematic of the representational similarity analysis. The same voxel-wise brain data and model activations used in the regression analysis were analyzed using representational similarity analysis. Representational dissimilarity matrices (RDMs) were computed for the 165 sounds using both the neural responses and the model activations, and the resulting RDMs were then correlated to quantify representational correspondence. C) Variance in voxel responses explained by model representations of the same set of 165 natural sounds. Brain data is from a previous fMRI study [51]. Graphs plot variance explained by best-predicting model stage, selected using held-out data (see Methods). Here and in C, analysis was performed on all sound responsive voxels in the auditory cortex whose responses to natural sounds exceeded a reliability threshold. Upper and lower horizontal lines plot variance explained by spectrotemporal and cochleagram baseline models’ respectively. Error bars plot SEM, derived by bootstrapping across sounds. D) Correlation between representational dissimilarity matrices of model representations and brain representations of the same set of 165 natural sounds (same brain data set as used for panel C). Graph plots correlation for most similar model stage, selected using held-out data (see Methods). Upper and lower horizontal lines plot RDM similarity for spectrotemporal and cochleagram baseline models’ respectively. Error bars plot SEM, derived by bootstrapping across sounds. E) Comparison of human-model brain alignment to human-model behavioral alignment. F) Comparison of model recognition performance (as measured by average d’ across benchmark conditions in each of the two experiments) to human-model brain alignment.

All optimized models demonstrated better regression-based brain predictions than the baseline models and the permuted controls, as in prior work with other task-optimized models (Fig. 8a) [15,52]. However, the models pre-trained on large datasets (CochCNNPretrained, VGGishPretrained, and SSASTPretrained) yielded better predictions than the models just optimized on the EnvAudioScene dataset alone. The RSA analysis showed weaker overall similarity between the trained models and brain representations, as in prior work, and only one model exhibited greater similarity than the ST baseline (Fig. 10c). But the models pre-trained on large datasets again showed greater brain-model similarity.

Figure 10c and 10d compare human-model brain similarity to human-model behavioral similarity (10c) and a summary measure of model recognition performance (10d). Across the models tested here, there is a tendency for the models with higher brain similarity to have higher behavioral similarity, and higher overall performance. This effect is similar to effects seen previously in models of other auditory [15] or visual tasks [53], and is consistent with the idea that the demands of mediating recognition tasks cause some degree of convergence in the representations of recognition systems [54]. These results suggest that training models on more diverse datasets causes them to both perform better and to produce more human-like behavior and more brain-like representations, although the match to either human behavior and brain representations was far from perfect.

## Discussion

We established a large-scale behavioral benchmark for human environmental sound recognition (EnvAudioEval) and used it to evaluate a set of computational models. The benchmark includes a broad variety of natural sounds, audio distortions, and multi-source scenes. Humans exhibited consistent patterns of performance across sound categories, distortion types, and scene sizes. These behavioral results provide a foundation for model comparison and validation.

We evaluated several classes of computational models—including biologically inspired baselines, convolutional neural networks (CNNs), and a transformer-based architecture—on the same recognition tasks used with human listeners. Models with learned representations, particularly those trained on large-scale datasets, produced the best matches to human performance (both overall performance, and the pattern of performance across conditions). The models producing better alignment with human behavior also produced better alignment with human brain representations as measured with two common metrics. Overall, the results suggest that many aspects of human recognition behavior emerge in systems optimized for real-world auditory classification tasks, but also reveal shortcomings in human alignment (and robustness) for all the models we tested.

### Relation to prior work

One goal of this work was to create a comprehensive benchmark for environmental sound recognition, expanding on prior studies that used smaller stimulus sets [6,7,9]. Our benchmark experiments include substantially more sounds (2176) compared to previous studies, which used 70 [6] and 168 [7] sounds, for instance, and a substantially larger set of signal distortions (68 for our experiments vs. 20 in Gygi et al., 2004, for instance). The larger scale enables a more comprehensive test of auditory recognition. We also conducted the first large-scale measurement of human recognition performance in multi-source scenes, and the first large-scale measurement of human environmental sound confusions. The stimuli and associated human data will be made publicly available following publication.

We leveraged this behavioral benchmark to evaluate computational models of environmental sound recognition for their match to human behavior. While prior work has shown that models trained on large audio datasets can achieve high classification accuracy [11–13,55–58], their alignment with human environmental sound perception had not been extensively tested. Our findings show that such models achieve closer correspondence with human behavior compared to traditional auditory models, with models pre-trained on more diverse datasets showing the best overall match across conditions. This latter result suggests that improvements to training data (in scale, variation, and realism) may be one way to obtain better matches to human behavior. Going forward, larger data sets are likely to be possible by combining recorded scenes with those generated by simulators [59–63]. The results are generally consistent with the idea that human behavior reflects the constraints of having been optimized for natural sounds and tasks [15–19,64,65], in that models optimized for such sounds and tasks provide better matches to behavior than other classes of models.

The behavioral benchmark also enabled us to compare model-brain similarity, which has been fairly extensively measured for responses to natural sounds in prior studies [15,52,66–68], to model-human behavioral similarity and model task performance, which has not previously been measured for environmental sound recognition. The model-brain similarity analyses replicated previous results with other artificial neural network models, but additionally found that the models that exhibited better behavioral alignment with humans also exhibited better brain alignment. This pattern is consistent with recent findings that models capturing human similarity ratings for natural sounds also better predict auditory-cortical representations [68].

### Limitations

Our recognition benchmark was based on a classification task, in part to enable comparisons with contemporary machine hearing models that perform classification. However, this choice constrained the benchmark to a pre-defined set of sound categories, which does not fully capture human representations of environmental sounds. In particular, human sound representations are often hierarchical (for instance, the screech of wheels and the honk of a horn could also be labeled as the sound of a car), with different levels of description that capture acoustic properties, physical properties of the source, or higher-level semantic interpretations [11,69,70]. Human sound recognition is also not always expressed in terms of verbal categories. For instance, physical events are often recognizable in terms of some of the underlying physical object properties [69,71–75]. As a result, a model that fully matches human behavior in our benchmark might nonetheless fail to replicate the richness of human recognition, and one should not conclude from the present results that machine systems are fully solving sound recognition as well as humans.

Another limitation is that all models were trained or fine-tuned on the EnvAudioScene training dataset, which was in turn built from the GISE-51 dataset. At the time of this work this seemed the best available source of clean, high-quality, labeled recordings of individual sound sources (which was essential for generating scenes with combinations of sources that were fully labeled). These virtues come at the cost of being small compared to some other audio datasets: GISE-51 contains only 16,357 unique audio clips, whereas AudioSet comprises of over 2 million clips. This limited training diversity may have imposed an upper bound on model performance and generalization even when models were first trained on a larger dataset and then fine-tuned.

The work presented here shares a common limitation of modeling work based on machine learning. Although some of the models tested here exhibit good quantitative matches to human behavior, they do not provide traditional explanations of human perception in terms of specific mechanistic hypotheses. Instead, perceptual abilities are explained as consequences of optimization. A skeptic might argue that we have traded one system we do not understand (the brain) for another that we do not understand (an artificial neural network). In response, we would highlight three contributions. First, the results show that task optimization alone is sufficient to get within striking distance of human recognition abilities on the benchmark experiments. A priori it was plausible that this might not be the case. Second, a model that matches human behavior relatively well, even if the model is difficult to interpret, provides a powerful tool for science. Because experiments on models are inexpensive, the models can be used to screen large numbers of experimental conditions to search for interesting structure that can then be probed for in humans [19]. A model also provides a baseline with which humans can be compared in future studies. For instance, because the models were trained on randomly generated scenes, they provide a baseline reference for what might be expected in a recognition system that does not leverage the co-occurrence statistics of sound sources in auditory scenes, allowing a stronger test of whether humans leverage such regularities between commonly co-occurring sources in the world [76]. Third, the infrastructure established here allows other models to be compared to those here, helping to enable cumulative science. Finally, although we have not attempted to probe the inner workings of the models we tested, we note that future studies could in principle probe the strategies learned by the models in ways that allow aspects of them to be lesioned or otherwise modified, potentially helping to clarify the representational transformations that underlie recognition.

### Future directions

The EnvAudioEval behavioral benchmark introduced here provides a starting point for quantifying the extent to which models can account for human auditory recognition behavior. As assessed by the benchmark, none of the models we tested here were fully adequate as descriptions of human behavior, even in the narrow domain of labeled categorization. One important direction will be to explore whether models that can fully explain human categorization judgments can be obtained simply by applying deep-learning-based classification models to bigger and better audio data sets that will likely become available in coming years. Self-supervision might be one way to train on much larger datasets, and the benchmark provides a way to assess whether future models trained with self-supervision better match human perception. Self-supervision is also likely closer to how humans learn from experience over the course of their life, and thus might in principle yield more human-like representations for this reason as well. Another promising avenue is the use of generative audio models to synthesize large-scale training datasets, which could provide fine-grained control over sound class diversity in ways that are difficult to achieve through collection of natural recordings alone.

The EnvAudioEval benchmark itself could also be augmented to more thoroughly probe human perception. A more complete characterization of human auditory scene perception might also include comparisons of structured versus random environments, to quantify contextual influences on recognition [76,77]. Expanding the benchmark to include spatialized multi-source scenes would allow exploration of how spatial layout influences source recognition [78,79] and whether models trained to recognize sounds with binaural input can capture human robustness in complex auditory environments.

The experiments and models developed here could also be extended to explore new questions in auditory perception. For instance, the experimental task, when applied to auditory scenes, provides an objective measure of the salience of a sound source in that scene. The models could thus be viewed as models of salience [80–83] and could be leveraged to generate hypotheses regarding the most salient sound sources across a range of auditory scene contexts. As seen in Figure 2g, sound recognition is also likely to be influenced by attention [84]. Such influences could be measured by cueing participants to particular locations, or to particular sound categories, and measuring the effect on recognition at scale. Attentional cueing mechanisms can also be incorporated into models [19] to further help clarify whether recognition limits in auditory scenes (Fig. 2b) can be mitigated by attention.

## Methods

### Human behavioral benchmark data collection procedure

#### Ethics statement

All participants provided written informed consent and the Massachusetts Institute of Technology Committee on the Use of Humans as Experimental Subjects (COUHES) approved all experiments.

#### Online experiment platform

To enable reliable measurements of human performance across large sets of experimental conditions, experiments were conducted using an online data collection platform (Prolific) to facilitate large sample sizes. Our laboratory has repeatedly found that online data can be of comparable quality to data collected in the lab provided a few modest steps are taken to standardize sound presentation, encourage compliance and promote task engagement [15,22–28].

Prolific participants were required to be in the United Kingdom or the United States, to have an approval rate of greater than 85%, and to self-identify as fluent in English. Participants were asked to perform the experiment in a quiet location and minimize external sounds as much as possible. They were instructed to set the computer volume to a comfortable level while listening to a sample of pink noise that was rms normalized to 0.005.

#### Sample sizes

Target sample sizes for each experiment were determined via a power analysis designed to estimate the reliability of the primary outcome metric. For Experiment 1a, we plotted split-half reliability of the target outcome metric as a function of sample size for subsamples of the pilot data (100 subsamples for each subsample size, from which we plotted the 20^th^ percentile). We then extrapolated the plotted curve to estimate the sample size required to achieve a desired reliability criterion 80% of the time. The target metric was the split-half reliability of the matrix of correlations between category-wise performance for different scene sizes (Figure 2d), which we intended to exceed.9. This yielded a target sample size of 160 participants.

Experiment 1b was run as part of a study examining effects of spatial separation on sound recognition, and we used the data from the control condition in which sound sources were co-located. We analyzed the data from all 8 participants that were run in that study; there was no power analysis to determine the sample size.

For Experiment 1c, we recruited 100 participants. Sample size was determined using a reliability-based power analysis based on pilot data. Specifically, we computed the split-half reliability of the target outcome metric (performance vs. scene size for the cued and uncued conditions) across subsamples of varying sizes (100 subsamples per size), plotting the 20th percentile of reliability estimates as a function of sample size. We then extrapolated this curve to estimate the sample size that would achieve the desired reliability of r ≥.9.

For Experiment 2, we collected 200 participants. This sample size was determined based on the structure of the experimental design: with 200 participants, each off-diagonal cell of the confusion matrix would be represented by 8 trials, which we hoped would be enough to estimate confusion rates to some extent. We confirmed post hoc that there was enough data to have split-half reliability that was well above chance.

In Experiment 3, participants were recruited in sets of 34, corresponding to the number required to fully cover all experimental conditions. Because randomly sampling participants often resulted in uneven representation of conditions, we estimated reliability using sets of 34 participants that preserved the balanced structure of the dataset (using pilot data from an experiment with the same structure as the actual experiment). Specifically, we computed the reliability of the target metric with subsamples of two and four sets of 34. The target metric was the split-half reliability of the distortion-wise performance (Figure 3b), which we intended to exceed.9. By extrapolating, we estimated that six sets of 34 participants would be required to achieve the desired reliability. This yielded a target sample size of 204 participants. Following the initial round of peer review, we collected two additional sets of 34 participants (total N = 272) to address reviewer questions about non-monotonicities in a few of the results graphs (Figure 3c; these smoothed out with more data).

#### Recognition task

In each experiment, on each trial participants were presented with a sound stimulus, followed by a prompt asking “Was [sound-category] present in the previous scene?”. Participants responded “yes” or “no” based on whether they detected the prompted source category. The probed category was present in the stimulus on half of the trials.

#### Familiarization

At the beginning of each experiment, participants completed a brief familiarization procedure to become acquainted with the set of possible labels used in the task. Participants were presented with two successive example recordings for each sound label while the label was displayed on the screen. These exemplars were sampled from the validation set of the GISE-51[45] dataset and were not used in the main experiment. This allowed participants to hear representative exemplars of each category.

#### Exclusion criteria

Participants had to meet two criteria to be included in the analysis. First, each experiment began with a headphone check task [85] to help ensure participants were wearing headphones and to promote consistent sound presentation across individuals. Headphone use also minimizes the influence of environmental noise and thus helps to standardize listening conditions. Participants who failed more than one trial in the headphone check were excluded from the experiment. Second, the experiments included ten catch trials intended to screen out participants who were not fully engaged with the task. The stimuli for these ten trials were randomly sampled from a set of 50 selected by the experimenter to be clearly identifiable single-source sounds. The ten catch trials were randomly interspersed within the main experiment. Participants who scored below 70% accuracy on a set of ten catch trials were excluded from analysis.

#### Analysis

Performance was quantified as d’:

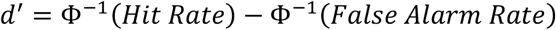

where Φ^−1^ is the inverse cumulative distribution function of the standard normal distribution. Hits were defined as affirmative responses (“yes”) when the probed source label corresponded to a source present in the auditory scene. False alarms were defined as affirmative responses when the probed label did not correspond to any source present in the scene.

### Experiment 1a: Environmental sound recognition in multi-source scenes - Online

#### Participants

250 participants were recruited through Prolific. Of these, 54 were excluded based on the exclusion criteria described above, yielding a final sample of 196 participants whose data were analyzed. Of these, 49% identified as female. The average age of participants was 39.88 years (s. d. = 13.56).

#### Procedure

Participants completed a total of 408 trials, split into four blocks of 102 trials. Within each block, every category was probed twice, with the probed category being present on one of those two trials. For each trial, the scene size was randomly selected from between 1 and 5 sources. Participants initiated each trial with a button click. The sound stimulus was then played once, followed by the query.

#### Stimuli

To test human participants on a diverse set of natural sounds using a stimulus set that was also suitable for model evaluation, we generated scenes using sounds from the evaluation portion of the GISE-51 dataset [45], as these were not included in model training. For all sounds longer than 1 second we excerpted the first second to preserve the sound onset. A 10-millisecond half-Hanning window was applied multiplicatively to the onset and offset of each sound. All stimuli were sampled at 44.1 kHz (the native sampling rate of the GISE dataset).

Scenes were generated with the constraint that no source category was used more than once within a scene. The non-probed categories for a scene were randomly selected subject to this constraint. Each of the sounds in the scene was normalized to the same rms level, and the waveforms were then summed. The onset of each sound in a scene was randomly jittered in time, uniformly sampled from between 0 and 1 second, resulting in scenes of up to 2 seconds. The final stimulus waveform for the scene was then rms normalized to 0.01.

### Experiment 1b: In-person replication of Experiment 1a

#### Participants

8 participants were recruited from the local community. Of these, 62.5% identified as female. The average age of participants was 24.38 years (s. d. = 7.42).

#### Procedure

Participants completed a total of 612 trials, split into six blocks of 102 trials. Scene size was randomly selected from between 2 and 5 sources rather than 1 and 5. Sounds were presented over a 133-speaker array, with scenes played from a randomly selected speaker. Participants responded using an iPad (same task as in Experiment 1a). On half of the trials the sounds in the scene were played from different speakers; these trials were excluded from the current analysis.

#### Stimuli

The experimental stimuli were identical to Experiment 1a, with the exception that sounds were presented at 60 dBA.

### Experiment 1c: Cueing manipulation

#### Participants

131 participants were recruited through Prolific. Of these, 23 were excluded based on the exclusion criteria described above, yielding a final sample of 108 participants whose data were analyzed. Of these, 48.15% identified as female. The average age of participants was 38.27 years (s. d. = 11.56).

#### Procedure

The experimental procedure was identical to Experiment 1a, with the exception that in two of the blocks, the participant was cued to the queried sound category label before the trial began. Participants were then presented the queried label again after the scene, identical to other scenes.

#### Stimuli

The experimental stimuli were identical to those of Experiment 1a.

### Experiment 2: Environmental sound recognition of single sources

#### Participants

246 participants were recruited through Prolific. Of these, 46 were excluded based on the exclusion criteria described above, yielding a final sample of 200 participants whose data were analyzed. Of these, 34% identified as female. The average age of participants was 33.6 years (s. d. = 9.88).

#### Procedure

Participants completed a total of 204 trials. Each participant was played 4 randomly selected exemplars from each sound category. On half the trials participants were probed with the correct label. On the remaining trials participants were probed with a different label, selected such that across sets of 25 participants, every label was probed once for every sound category presentation. Participants initiated each trial with a button click. The sound stimulus was then played once, followed by the query.

#### Stimuli

The same sounds and preprocessing procedures from Experiment 1a were used to obtain source signals. The resulting stimulus waveforms were truncated in time to a duration of one second, windowed at onset and offset with 10 ms half-Hanning windows, and normalized to have a root mean square (RMS) amplitude of 0.01.

### Experiment 3: Recognition of distorted environmental sounds

#### Participants

339 participants were recruited through Prolific. Of these, 67 were excluded based on the exclusion criteria described above, yielding a final sample of 272 participants whose data were analyzed. Of these, 54% identified as female. The average age of participants was 38.85 years (s. d. = 12.81).

#### Procedure

At the start of the experiment, participants were informed that the sounds could be distorted in various ways. The experiment included a total of 3,468 unique conditions, spanning all combinations of 68 distortions and 51 sound categories. Trials were generated for sets of 17 participants to enable all conditions to be sampled once across the set of 17 participants, with each participant completing 3,468/17 = 204 trials. The subsets of 204 trials for each participant used each of the 68 distortions 3 times and each of the 51 sound categories 4 times as the queried category. The probed category present on half of the trials (“target-present” trials); the half of the trials that had target-present status was randomly selected from all trials. The same assignment of distortions and sounds was used for two complementary sets of 17 participants with the target-present trials flipped for the two sets, such that every set of 34 participants yielded one target-present and one target-absent trial for each distortion x category combination. Participants initiated each trial with a button click. The sound stimulus was then played once, followed by the query.

#### Stimuli

The same sounds and preprocessing procedures from Experiment 1a were used to obtain source signals to which distortions were applied. Distortions were applied as described in Table 1. The resulting stimulus waveforms were truncated in time to a duration of one second, windowed at onset and offset with 10 ms half-Hanning windows, and normalized to have a root mean square (RMS) amplitude of 0.01.

**Table 1.** Sound distortion and implementation details. This table summarizes the acoustic distortions applied to the stimuli (Condition), the specific distortion levels used for each Condition (Distortion Levels), and the procedures used to generate or apply each distortion (Procedure).

| Condition | Distortion Levels | Procedure |  |
| --- | --- | --- | --- |
| Original |  |  | Baseline for comparison; establishes ceiling performance across listeners and tasks. |
| High-pass Filter | Cutoffs: 400, 800, 1600, 3200, 6400 Hz | A fourth-order high-pass Butterworth filter was applied in both forward and reverse directions to avoid phase shifts. | Reveals importance of low frequency information and invariance to spectral filtering. |
| Low-pass Filter | Cutoffs: 400, 800, 1600, 3200, 6400 Hz |  | Reveals importance of high frequency information and invariance to spectral filtering. |
| Band-pass Filter with varied Center Frequency | Cutoffs (Hz): [150, 300], [300, 600], [600, 1200], [1200, 2400], [2400, 4800], [4800, 9600] |  | Reveals the perceptual weighting of each frequency band, importance of spectral context, and invariance to spectral filtering. |
| Band-pass Filter with varied Bandwidth | Cutoffs (Hz): [1124, 2002], [842, 2673], [472, 4762], [265, 8485], [149, 15119] |  |  |
| Time Compression / Dilation | Tempo change factor: 0.5, 0.75, 0.875, 1.125, 1.25, 1.5, 2 | The tempo of each signal was altered using the Waveform Similarity Overlap-Add (WSOLA) algorithm as implemented in Sox [86], which preserves pitch while modifying temporal structure. Tempo change factor specifies tempo as proportion of original tempo. | Reveals the importance of temporal structure and invariance to tempo. |
| Reverberation | Direct-to-Reverberant (DRR): 20, 50, 80 dB<br>x<br>RT60: 200, 400, 800, 1600 ms | The signal was convolved with synthesized impulse responses, following the method of Traer and McDermott (2016) [33]. Impulse responses were synthesized by passing pink noise through a filterbank approximating human cochlear processing. Then the resulting subbands were modulated by exponentially decaying envelopes (with frequency-dependent decay constants modeled after real-world environmental impulse responses) before being summed to produce the final impulse response. Three | Simulates real-world listening conditions, testing robustness to distortion imposed by natural acoustic environments. |
|  |  | DRR conditions were crossed with four RT60 conditions. |  |
| Local Time Reversal | Segment lengths: 10, 20, 30, 40, 50, 100 ms | Each sound was divided into adjacent non-overlapping segments. Each segment was then time-reversed, and the reversed segments were concatenated to yield the final signal. | Reveals the importance of fine-grained temporal structure. |
| Peak Clip | Threshold (percentile): 0, 0.25, 0.5, 0.75, 0.9, 0.98 | All samples in the signal that were above the top nth percentile for that signal were clipped to that threshold. | Reveals the auditory system's tolerance for waveform distortion, by reducing dynamic range and introducing harmonic distortion. |
| Noise Vocoding | Number of channels: 1, 2, 4, 8, 16, 32 | Signals were filtered with ERB-spaced filters. The resulting subbands were grouped into channels and summed (this was a simple way to generate different sets of channels that each tiles the frequency spectrum). Hilbert envelopes were extracted for each channel (providing "target" envelopes used for noise vocoding). Gaussian noise was filtered through the same channels, its envelopes replaced with the sound envelopes (by dividing out the noise envelope and multiplying by the target envelopes), and the outputs summed to produce the final waveform. | Reveals dependence on spectrally resolved information. |
| Spectral Lowpass Modulation Filtering | 0.5, 1, 2, 4, 8 cycles/kHz | A spectral lowpass filter was applied in the modulation domain, attenuating modulations above a specified cutoff frequency before reconstructing the filtered signal. Modulation filtering was performed using the method of Elliot and Theunissen (2009) [30], using code supplied by Frederic Theunissen. | Reveals dependence on spectral detail at different resolutions. |
| Temporal Lowpass Modulation Filtering | 3, 6, 12, 24 Hz | A temporal lowpass filter was applied in the modulation domain, attenuating temporal modulations above a specified cutoff frequency before reconstructing the filtered signal. Modulation filtering was performed using the method of Elliot and Theunissen (2009) [30], using code supplied by Frederic Theunissen. | Reveals dependence on temporal detail at different resolutions. |

### Model training

#### EnvAudioScene training data set

We generated training data using source signals from the GISE-51 training dataset [45]. For each training example, we first sampled a scene size (i.e., the number of concurrent sources) uniformly from one to five. Sound source categories and exemplars were then selected at random for each scene, subject to the constraint that no category was used more than once per scene. Each individual source signal for the sampled category and exemplar was truncated or zero-padded to a duration of one second, with a 10 ms half-Hanning window applied at both the onset and offset. To introduce temporal variability, the onset of each source signal was randomly sampled from 0 to 1 seconds. The full scene duration was fixed at two seconds. We generated a total of 1.5 million auditory scenes for the training dataset.

Scenes were rendered using a spatialization procedure to create realistic and varied reverberation (even though models were monaural). Spatialization followed the procedure of Francl and McDermott (2022) [17], using the rooms generated by Saddler and McDermott [16]. For each example, one of 2,000 acoustically modeled rooms was chosen at random. Within the selected room, each source was assigned a random azimuth (uniformly sampled from 0–355° in 5° steps) and elevation (uniformly sampled from 0–60° in 10° steps) from the listener’s position, yielding 504 potential source directions. Source distances were drawn uniformly from 1.4 to 29 meters. Source signals were zero-padded to 2.5s before spatialization. The spatialization procedure produced a binaural waveform; because the models were monaural we took the left channel of the spatialized audio for the training data.

#### AudioSet pretraining

For models pretrained on AudioSet [44], we used the strongly labeled subset of AudioSet as training data (i.e., for which the labels were time-indexed). Sound clips containing less than 0.5 seconds of labeled content were excluded. For each remaining clip, a random two-second segment containing labeled sound was selected, and all labels present during that segment were included in the training target labels. To add reverberation and increase variability, each selected clip was spatialized to a randomly chosen location within a virtual room using the procedure described in the previous section.

#### Training procedure

All models performed a multi-label classification task. Each model had independent sigmoidal output classifier units for each sound category, trained using binary cross-entropy loss (with one term in the loss for each category) with a learning rate of 1e-4. Optimization was performed using the Adam optimizer with β₁ = 0.9, β₂ = 0.999, and ε = 1e-7. Training was conducted with a batch size of 32. To ensure comparability, all models were trained until they reached asymptotic performance as determined by the early stopping function in PyTorch, which stopped training if loss had not dropped for 3 validation cycles (with each cycle comprising a pass through a tenth of the training data). Training was performed on the Open Mind computing cluster at MIT using NVIDIA A100 GPUs over about four days.

Models pretrained on AudioSet were pretrained prior to finetuning on the EnvAudioScene dataset/task. Pretraining used the early stopping procedure described above, which always terminated training within 5 epochs. Baseline and pretrained models were then fine-tuned on the EnvAudioScene dataset (for 1 epoch). Fully optimized CNN models were trained using the early stopping procedure, which always terminated training within 2 epochs (we found that models tended to overfit if trained beyond this point).

### Cochleagram generation

Cochleagram generation was similar to previous papers in our lab, and the methods description is reproduced from one recent paper [19] with minor edits.

#### Overview

The first stage of our models was a fixed simulation of the cochlea and auditory nerve, providing a time-by-frequency representation intended to replicate the auditory cues provided by the human auditory periphery. This initial stage took the sound waveform as input, sampled at 48 kHz. The input was passed through a finite-impulse-response (FIR) approximation of a gammatone filter bank (with impulse responses truncated to 25 ms to reduce memory consumption) the output of which was half-wave rectified, passed through a compressive nonlinearity, then downsampled.

#### Implementation details

First, the 48 kHz audio waveform was convolved in the time domain with the FIR approximation of a 211-channel gammatone filter bank with center frequencies spaced uniformly on an ERB-numbered scale between 40Hz to 24kHz. Second, the resulting subbands were half-wave rectified. Third, the half-wave rectified subbands were raised to the power of 0.3 to simulate the compressive response mediated by outer hair cells. Fourth, the compressed, rectified subbands were low-pass filtered with a 4kHz cutoff and downsampled to 8kHz, to both impose the upper limit on phase locking of inner haircells and reduce the dimensionality of the neural network inputs. Low-pass filtering and downsampling was performed via 1d convolution with a Kaiser-windowed sinc filter, with a filter width of 64, roll-off of 0.94759, and beta of 14.76965, implemented with the Torchaudio transforms resample method. Finally, to avoid signal onset/offset artifacts, the middle 2-seconds were then excerpted from the full 2.5-second input signal duration resulting from the spatialization procedure described above. The resulting 211-frequency by 16,000-timestep representation, which we refer to as a “cochleagram”, was the input to the neural networks.

### Model architectures

#### Cochleagram baseline

To establish a baseline representing the performance of a model relying exclusively on the feature output from the auditory periphery, we trained a single-layer classifier operating on a 211-channel cochleagram. This model consisted of a linear readout layer that received the flattened cochleagram as input and produced sigmoid outputs for the 51 sound classes, analogous to the output layer used in the CNN architectures.

#### Spectrotemporal baseline

To establish a baseline representing the performance of a model relying on the feature output from a hand-engineered model of the auditory cortex, we trained a single-layer classifier operating on a SpectroTemporal modulation filter model based on Chi et al. (2005) [46]. The model consisted of two fixed stages preceding the classifier. The first stage was a cochleagram. The cochleagram was similar to that described above except that the representation was downsampled to 200 Hz and cropped to the central 1.95 seconds (to accommodate the dimensions expected by the instantiation of the modulation filter bank that we used). The second stage was a modulation filter bank.

The modulation filter bank architecture was similar to that used in previous papers from our lab, and the methods description here is reproduced from one recent paper with minor edits [52]. The main difference in our implementation relative to the original implementation of Chi et al. was that spectral filters were specified in cycles/erb (rather than cycles/octave) because the input signal to the model was a cochleagram generated with ERB-spaced filters. The model consisted of a linear filter bank tuned to spectrotemporal modulations at different frequencies, spectral scales, and temporal rates. The spectrotemporal filters were applied via a 2D convolution with zero padding in frequency (211 samples) and time (390 samples). Spectrotemporal filters were constructed with center frequencies for the spectral modulations of [0.0625, 0.125, 0.25, 0.5, 1, 2] cycles/erb, and center frequencies for the temporal modulations of [0.5, 1, 2, 4, 8, 16, 32, 64]. Both upward and downward frequency modulations were included (resulting in 96 filters). An additional 6 purely spectral and 8 purely temporal modulation filters were included for a total of 110 modulation filters. To extract the power in each frequency band for each filter, we squared the output of each filter response and took the average across time for each audio frequency channel, similar to previous studies [15,46,48,52]. We then trained a single linear readout layer on top of this time-averaged power representation to obtain sound category predictions. The readout had one sigmoid unit per sound category.

#### CochCNN

This architecture was adapted from prior work in our lab, which identified it through a systematic architecture search [16] to identify model architectures that produced good performance on word and voice recognition tasks using stimuli of the same length as used here. The input to the model was a cochleagram representation of the input sound, treated as a single-channel time–frequency image. The network architecture consisted of six convolutional blocks, each comprising a sequence of layer normalization, convolution, ReLU activation, and Hann-window pooling operations [52,87]. Hann window pooling was used to avoid aliasing artifacts associated with other types of pooling operations (e.g. max pooling) [87,88].

The feature map resulting from this convolutional backbone was flattened into a one-dimensional vector and passed through two fully connected layers. The first fully connected layer projected the flattened features into a lower-dimensional embedding space and was followed by layer normalization, dropout (with a dropout rate of 0.5), and a ReLU activation. The second fully connected layer produced a 51-dimensional output corresponding to the 51 sound classes. Each output unit applied a sigmoid nonlinearity to support multi-label classification.

#### CochVGGish

To evaluate the performance of a widely used externally developed architecture, we implemented a variant of the VGGish model. The architecture was identical to the original VGGish model, except that the neural network received cochleagrams as input rather than log-mel spectrograms. To align with the parameters of the original VGGish model, the cochleagram was downsampled to a temporal sampling rate of 100 Hz (note that this input temporal resolution is much lower than that of the in-house models). In addition, because the original VGGish architecture operated on 0.96-second inputs, we cropped the middle 0.96 seconds of each stimulus to generate model inputs. The model was trained from scratch on the GISE-51 dataset, using the same training procedure as the other task-optimized models described above.

#### CochSTVGGish

This model combined the spectrotemporal filterbank representation with a convolutional neural network. The first two stages were identical to the spectrotemporal filterbank model described above. The output of the filterbank was then passed to a CNN with a variant of the VGGish architecture used in the CochVGGish model. We selected the VGGish architecture for this hybrid model because the dimensionality and structure of the spectrotemporal model’s output closely matched the input format expected by VGGish. The neural network architecture was the same as that used in the original VGGish model except that the number of input frequency channels was increased from 64 to 211, the input length was doubled to accommodate 2 second sound signals, and the first fully connected layer was larger to accommodate the resultingly larger dimensions of the final convolutional stage.

#### CochCNNPretrained

This model had the same architecture as the CochCNN model, but was pretrained on the AudioSet training set and task, as described as described in the “Training procedure” section above. Following pretraining, we fine-tuned the model for 1 epoch on the EnvAudioScene dataset and task. During finetuning, all model parameters were updated (i.e., no weights were frozen). The only architectural modification during finetuning was to adjust the final fully connected layer to produce 51 output classes, rather than the 456 outputs used in the AudioSet task (and thus in the pretrained model), to match the labels in our dataset.

#### VGGishPretrained

This model was initialized from the trained PyTorch implementation of the standard VGGish network [11]. To adapt it to our task we appended a fully connected layer to map the model’s 128-dimensional embedding to the 51 output classes. Because the original VGGish architecture operates on 0.96-second inputs, we cropped the middle 0.96 seconds of each stimulus to generate model inputs. Given the way the stimuli are constructed, this preserved a large portion of each sound source, but could have limited model performance relative to that of humans. We maintained the original log-mel spectrogram input stage. The model was fine-tuned on the EnvAudioScene dataset without freezing any layers.

#### SSASTPretrained

This model was based on a self-supervised audio spectrogram transformer (SSAST) model [13] (we used the Hugging Face implementation, loading the pretrained SSAST weights into the AST model architecture). The model input consisted of log-Mel spectrograms as in the original SSAST implementation. To adapt the network to our task, the final dense layer was modified to 51 units corresponding to GISE-51 sound classes. The model was then finetuned on the EnvAudioScene dataset, with all parameters updated during training.

### Evaluating models on benchmark

#### Experiment 1 procedure

Each model was tested on the same set of multi-source sound scenes presented to human participants. Stimuli were upsampled to a sampling rate of 48 kHz to match the model’s training data. All inputs were normalized to match the level used during model training.

#### Experiment 2 procedure

Each model was tested on the same set of exemplars that were presented to human participants. Stimuli were up-sampled to 48 kHz and padded with 500 ms of silence on either side to conform with the model input requirements (because the individual sound source excerpts were 1s in duration). All inputs were normalized to match the level used during model training.

#### Experiment 3 procedure

Each model was tested on the same set of distorted environmental sounds presented to human participants. Stimuli were up-sampled to 48 kHz and padded with 500 ms of silence on either side to conform with the model input requirements (because the individual sound source excerpts were 1s in duration). All inputs were normalized to match the level used during model training.

#### Analysis

Because the models output class probabilities rather than binary judgments, we derived d′ values from the area under the receiver operating characteristic curve (AUC). The AUC was computed from the empirical cumulative distribution of model outputs for a given category, across all test scenes. This approach provides a criterion-free estimate of sensitivity that is comparable across models and experimental conditions. The alternative would be to impose a fixed decision criterion to generate discrete responses, but it is not obvious how to equate such a criterion across models.

To generate model confusion matrices for Experiment 2, we derived individual detection thresholds for each model and sound category. Using the experiment trials for each category, we identified the lowest possible threshold value that yielded a hit rate at or below the mean human hit rate in models, and then used this threshold to determine model responses.

### Voxel response modeling

Model-brain fMRI analysis was similar to that described in previous papers in our lab. The analysis used the code from the repository of one recent paper [52], and the methods description is reproduced from that paper with minor edits. This recent study [52] analyzed two different fMRI datasets [51,89] and obtained very similar results, and so here we restricted analysis to the earlier of the two datasets. We also included a Cochleagram baseline model in addition to the SpectroTemporal baseline model.

We note that the cochleagram representation used in the models in this study had a higher upper frequency limit than the cochleagram representations of the models in the previous study [52] (in part because we plan to extend the models to use spatial cues, and wanted to support the full range of head-related transfer function cues available to humans). This difference likely explains the quantitative differences in model-brain alignment metrics for the models that are common to both studies (e.g. the spectrotemporal model).

#### General

FMRI data was measured from the auditory cortex of humans listening to a set of 165 two-second natural sounds [51]. Sounds were presented in blocks of 5 repeated presentations of the same sound. One of the sounds was lower in intensity than the others, and the participants detected the intensity change, a task intended to force them to listen to the sounds. Responses were analyzed for all voxels that responded significantly more to sound than to silence, and whose response to natural sounds exceeded a threshold level of reliability.

We performed an encoding analysis in which each voxel’s time-averaged activity was predicted by a regularized linear function of the model activations in response to the same set of sounds used in the fMRI experiment. We operationalized each model stage within each candidate model as a hypothesis of a neural implementation of auditory processing. The fMRI hemodynamic signal to which we were comparing the candidate model blurs the temporal variation of the cortical response, such that the analysis was based on the time-average of the fMRI response to each sound (because there is little temporal structure in the fMRI response to a two-second sound). To compare the model activations to the fMRI data, we predicted each voxel’s time-averaged response to each sound from time-averaged model responses (obtained by averaging the model responses over the temporal dimension after extraction). Because it seemed possible that units with real-valued activations might average out to near-zero values, we extracted unit activations after ReLU stages. Transformer architectures had no such stages, so we extracted the real-valued unit activations and analyzed all model stages in this way.

#### Voxelwise modeling: Regularized linear regression and cross-validation

We modeled each voxel’s time-averaged response as a linear combination of a model stage’s time-averaged unit responses. We first generated 10 randomly selected train/test splits of the 165 sound stimuli into 83 training sounds and 82 testing sounds. For each split, we estimated a linear map from model units to voxels on the 83 training stimuli and evaluated the quality of the prediction using the remaining 82 testing sounds (described below in greater detail). For each voxel-stage pair, we took the median of the prediction quality across the 10 splits. The linear map was estimated using regularized linear regression. Given that the number of regressors (i.e., time-averaged model units) typically exceeded the number of sounds used for estimation (83), regularization was critical. We used L2-regularized (“ridge”) regression, which can be seen as placing a zero-mean Gaussian prior on the regression coefficients. Introducing the L2-penalty on the weights results in a closed-form solution to the regression problem, which is similar to the ordinary least-squares regression normal equation:

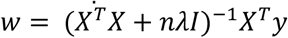

where w is a d-length column vector (the number of regressors—i.e., the number of time-averaged units for the given stage), y is an n-length column vector containing the voxel’s mean response to each sound (length 83), X is a matrix of regressors (n stimuli by d regressors), n is the number of stimuli used for estimation (83), and I is the identity matrix (d by d). We demeaned each column of the regressor matrix (i.e., each model unit’s response to each sound), but we did not normalize the columns to have unit norm. Similarly, we demeaned the target vector (i.e., the voxel’s response to each sound). By not constraining the norm of each column to be one, we implemented ridge regression with a non-isotropic prior on each unit’s learned coefficient. Under such a prior, units with larger norm were expected a priori to contribute more to the voxel predictions. The paper on which these methods were based [52] found that this procedure led to more accurate and stable predictions in left-out data, compared with a procedure where the columns of the regressor matrices were normalized (i.e., with an isotropic prior). The demeaning was performed on the train set, and the same transformation was applied on the test set. This ensured independence (no data leakage) between the train and test sets.

Performing ridge regression requires selecting a regularization parameter that trades off between the fit to the (training) data and the penalty for weights with high coefficients. To select this regularization parameter, we used leave-one-out cross-validation within the set of 83 training sounds. Specifically, for each of 100 logarithmically spaced regularization parameter values (1e-50, 1e-49,…,1e48, 1e49), we measured the squared error in the resulting prediction of the left-out sound using regression weights derived from the other sounds in the training split. We computed the average of this error (across the 83 training sounds) for each of the 100 potential regularization parameter values. We then selected the regularization parameter that minimized this mean squared error. Finally, with the regularization parameter selected, we used all 83 training sounds to estimate a single linear mapping from a stage’s features to a given voxel’s response. We then used this linear mapping to predict the response to the left-out 82 test sounds and evaluated the Pearson correlation of the predicted voxel response with the observed voxel response. If the predicted voxel response had a standard deviation of exactly 0 (no variance of the prediction across test sounds), the Pearson correlation coefficient was set to 0. Similarly, if the Pearson correlation coefficient was negative, indicating that the held-out test sounds were not meaningfully predicted by the linear map from the training set, the Pearson correlation value was similarly set to 0. We squared this Pearson correlation coefficient to yield a measure of variance explained. As in the paper on which these methods were based [52], we found that the selected regularization parameter values rarely fell on the boundaries of the search grid, suggesting that the range of the search grid was appropriate. We emphasize that the 82 test sounds on which predictions were ultimately evaluated were not incorporated into the procedure for selecting the regularization parameter nor for estimating the linear mapping from stage features to a voxel’s response—i.e., the procedure was fully cross-validated.

#### Voxelwise modeling: Correcting for reliability of the measured voxel response

The use of explained variance as a metric for model evaluation is inevitably limited by measurement noise. To correct for the effects of measurement noise, we computed the reliability of both the measured voxel response and the predicted voxel response. Correcting for the reliability of the measured response is important to make comparisons across different voxels, because (as shown in, for instance, Figure S2 in Kell and colleagues’ article [31]) the reliability of the BOLD response varies across voxels. This variation can occur for a variety of reasons (for instance, distance from the head coil elements). Not correcting for the reliability of the measured response will downwardly bias the estimates of variance explained and will do so differentially across voxels. This differential downward bias could lead to incorrect inferences about how well a given set of model features explains the response of voxels in different parts of auditory cortex.

#### Voxelwise modeling: Correcting for reliability of the predicted voxel response

Measurement noise corrupts the test data to which model predictions are compared (which we accounted for by correcting for the reliability of the measured voxel response, as described below), but noise is also present in the training data and thus also inevitably corrupts the estimates of the regression weights mapping from model features to a given voxel. This second influence of measurement noise can be addressed by correcting for the reliability of the predicted response. Doing so is important for two reasons. First, as with the reliability of the measured voxel response, not correcting for the predicted voxel response can yield incorrect inferences about how well a model explains different voxels. Second, the reliability of the predicted response for a given voxel can vary across feature sets, and failing to account for these differences can lead to incorrect inferences about which set of features best explains that voxel’s response. By correcting for both the reliability of the measured voxel response and the reliability of the predicted response, the ceiling of our measured r-squared values was 1 for all voxels and all stages, enabling comparisons of voxel predictions across models.

#### Voxelwise modeling: Corrected measure of variance explained

To correct for the reliability of the measured and predicted voxel responses, we employed the correction for attenuation [90]. This correction is a standard technique in many fields and is becoming more common in neuroscience. The correction estimates the correlation between two variables (here the measured voxel response and the model prediction of that response) independent of measurement noise. The result is an unbiased estimator of the correlation coefficient that would be observed from noiseless data. Our corrected measure of variance explained was the following:

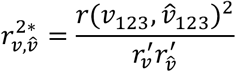

where *v*_123_ is the voxel response to the 82 left-out sounds averaged over the 3 scans, *v̂*_123_ is the predicted response to the 82 left-out sounds (with regression weights learned from the other 83 sounds), r is a function that computes the correlation coefficient, *r’_v_* is the estimated reliability of that voxel’s response to the 83 sounds, and *r’_v̂_* is the estimated reliability of that predicted voxel’s response. *r’_v_* is the median of the correlation between all 3 pairs of scans (scan 0 with scan 1; scan 1 with scan 2; and scan 0 with scan 2), which is then Spearman–Brown corrected to account for the increased reliability that would be expected from tripling the amount of data [90]. *r’_v̂_* is analogously computed by taking the median of the correlations for all pairs of predicted responses (models fitted on a single scan) and Spearman–Brown correcting this measure. Note that for very noisy voxels, this division by the estimated reliability can be unstable and can cause for corrected variance explained measures that exceed one. To ameliorate this problem, we limited both the reliability of the prediction and the reliability of the voxel response to be greater than some value *k* [91]. For *k* = 1, the denominator would be constrained to always equal 1, and, thus, the “corrected” variance explained measured would be identical to the uncorrected value. For *k* = 0, the corrected estimated variance explained measure is unaffected by the value *k*. This k-correction can be seen through the lens of a bias-variance trade-off: the correction reduces the amount of variance in the estimate of variance explained across different splits of stimuli but does it at the expense of a downward bias of those variance-explained metrics (by inflating the reliability measure for unreliable voxels). For *r’_v_*, we used a k of 0.182, which is the *p* < 0.05 significance threshold for the correlation of two 83-dimensional Gaussian variables (i.e., with the same length as our 83-dimensional voxel response vectors used as the training set), while for *r’_v̂_*, we used a k of 0.183, which is the *p* < 0.05 significance threshold for the correlation of two 82-dimensional Gaussian variables (i.e., same length as our 82-dimensional predicted voxel response vectors, the test set).

#### Voxelwise modeling: Aggregate measures

We repeated the explained-variance estimation procedure for each stage and voxel 10 times, once each for 10 random train/test splits, and took the median explained variance across the 10 splits for a given stage-voxel pair. We performed this procedure for all stages of all candidate models and all voxels (7,694 voxels). Thus, for each stage and voxel, this resulted in 10 explained variance values (R^2^). We computed the median explained variance across these 10 cross-validation splits for each voxel-stage pair. For comparison, we performed an identical procedure with the stages of models with permuted weights (with the same architecture as the trained models) and the Cochleagram and SpectroTemporal baseline model. In all analyses, if a noise-corrected median explained variance value exceeded 1, we set the value to 1 to avoid an inflation of the explained variance.

To compare how well each candidate model explained the variance in the fMRI data, we aggregated the explained variance across all voxels for each model. We evaluated each candidate model using its best-predicting stage. For each voxel, we used half of the 10 cross-validation test splits to select the best-predicting stage and the remaining 5 test splits to obtain the median explained variance. This yielded a median explained variance per voxel. To ensure that this procedure did not depend on the random 5 cross-validation splits selected, we repeated this procedure 10 times for each model. We then obtained the mean of the explained variance values for each voxel across these 10 iterations. To obtain the aggregated explained variance across voxels for each model, we first obtained the median across voxels within each participant and then took the mean across participants. An identical procedure was used for the permuted networks.

#### Model-brain representational similarity analysis

Representational similarity analysis (RSA) was identical to the analysis in a recent paper from our lab [52], with the exception of adding the cochleagram baseline. The methods description in the following paragraph is reproduced from that paper [52] with minor edits.

In the analysis shown in Figure 8b, we compared the representational dissimilarity matrices (RDMs) computed across all voxels of a human participant to the RDM computed from the time-averaged unit responses of each stage of each model in response to each of the sounds used in the fMRI experiment. To choose the best-matching stage, we first generated 10 randomly selected train/test splits of the 165 sound stimuli into 83 training sounds and 82 testing sounds. For each split, we computed the RDMs for each model stage and for each participant’s fMRI data for the 83 training sounds. We then chose the model stage that yielded the highest Spearman ρ measured between the model stage RDM and the participant’s fMRI RDM. Using this model stage, we measured the model and fMRI RDMs from the test sounds and computed the Spearman ρ. We repeated this procedure 10 times, once each for 10 random train/test splits, and took the median Spearman ρ across the 10 splits. We performed this procedure for all candidate models and all participants (8 participants, from a previously published fMRI study [51]) and computed the mean Spearman ρ across participants for each model. For comparison, we performed an identical procedure with permuted versions of each neural network model and with the Cochleagram and SpectroTemporal baseline model.

### Statistics

The reliability of human performance in Experiment 1a was estimated using split-half correlations across odd-and even-numbered participants (based on the order in which they were run). For the analysis involving correlations between category-wise performance for different scene sizes, the likelihood that the correlations involving a scene size of one could have been lowest just by chance was evaluated using an exact permutation test (there are 210 ways to choose 4 correlations out of the 10 total, only one of which yields the four correlations involving a scene size of one). In Experiment 1c the effect of cueing at a scene size of 1 was evaluated using a t-test and the effect of cueing across all scenes 1c was evaluated using a repeated measures ANOVA.

The reliability of confusions and of exemplar hit rates in Experiment 2 was estimated using split-half correlations across a split of the first and second half of participants to ensure even distribution of conditions across split (based on the order in which they were run). The effect of sound duration was tested using a repeated measures ANOVA.

To calculate the reliability of performance in experiment 3 we divided participants into splits with off-and even-numbered subsets of 34 participants (each subset of 34 participants covered all experimental conditions; subsets were run consecutively) and calculated split-half reliability between these groups. Human–model correspondence was assessed by comparing average d’ values for either sound categories or distortions, depending on the experiment, with each of two metrics: the Spearman correlation, and the root-mean-squared error.

The noise ceiling on correlations was estimated as the geometric mean of the reliabilities of the quantities being correlated. For model performance, reliability was estimated from splits of trials. For both humans and models, the split-half reliability was Spearman-Brown corrected [92] to estimate the reliability with the full set of participants and trials, respectively.

Model robustness to sound distortions was evaluated with a paired t-test comparing human and model performance.

Error bars reflect the standard error of the mean (SEM) from 1000 bootstrap resamples.

## Data and code availability

Data and code are available in a repository that will be made public at the time of publication. For the time being they are provided for editors and reviewers via Google drive: https://drive.google.com/file/d/1kmemz1ALKHtv2TYkUIczfZA6p5vWqjSM/view?usp=sharing

## Author contributions

S.A. designed and ran the human experiments, trained the in-house models, tested all models, made the figures, and wrote the paper. J.H.M. supervised the research, helped design the experiments, raised the funding for the research, and wrote the paper.

## Acknowledgements

Research supported by National Institutes of Health grants R01 DC017970 and R01 DC021464, and by a Friends of the McGovern Institute fellowship to S.A. The authors thank Annika Magaro for help with the selection and implementation of the sound distortions, and the McDermott lab and Dan Ellis for helpful feedback on the manuscript.

## Notes

### Competing Interest Statement

The authors have declared no competing interest.

### Summary of Updates

We added three new experiments to strengthen the manuscript. First, we included two additional control experiments to evaluate human performance in person and with semantic attention, providing further validation of the proposed approach. Second, we added an experiment that systematically analyzes model confusions, offering additional insights into the human's and model's behavior.

